# Equation-Based Integration of Flux Balance Analysis with Diffusion for Spatio-Temporal Simulation of Microbial Communities

**DOI:** 10.64898/2026.04.11.717857

**Authors:** Frederick Senya, Ryan Siegel, llija Dukovski, David Bernstein

## Abstract

Spatio-temporal interactions shape microbial community dynamics. Metabolism, through competition and cross-feeding, is a foundational mechanism of these interactions. Flux balance analysis enables efficient simulation of steady-state metabolism. Integrating these simulations through time, using dynamic flux balance analysis, provides temporal predictions of growth and metabolism. Incorporating spatial context, through partial differential equations, enables spatio-temporal simulation of microbial communities. In this chapter, we step through this sequential process, moving from steady-state, to temporal, to spatio-temporal simulation of microbial community metabolism. We provide an illustrative example using the modeling software COMETS (Computation of Microbial Ecosystems in Time and Space) to simulate interacting bacterial colonies of *Bifidobacterium longum* subsp. *infantis* and *Anaerobutyricum hallii* (previously *Eubacterium hallii*). Within this simulation, both competition and cross-feeding influenced the production of butyrate leading to an intermediate optimal interaction distance for metabolite production. We outline each step and provide open-source code such that this simulation can serve as a template for future spatio-temporal simulations of microbial community metabolism.

## 1 Introduction

Microbial organisms exist in complex, spatially-organized communities, where competition for nutrients and cross-feeding of metabolic byproducts define community structure and growth dynamics (Tropini et al., 2017; Bäcker et al., 2026; Dal Co et al., 2023; Mark Welch et al., 2016; Pignon and Schaerli, 2025; Kim et al., 2008). Computational simulation, with genome-scale metabolic models (GEMs), provides quantitative insight into the growth and metabolic dynamics of microbial communities (Quinn-Bohmann et al., 2025; Heinken et al., 2021b; Tarzi et al., 2024; Ankrah et al., 2021; Biggs et al., 2015; Heinken et al., 2021a). Simulating microbial community metabolism in space and time is a multi-scale approach that integrates efficient steady-state simulations of metabolism with dynamic simulations of the propagation of biomass and metabolites (Dukovski et al., 2021; Harcombe et al., 2014; Bauer et al., 2017; Biggs and Papin, 2013; Cole et al., 2015; Chen et al., 2016; Borer et al., 2019). Within these multi-scale simulations, individual organisms optimize their own growth and consume/produce metabolites in their local environments. These local environmental changes diffuse throughout the system, impacting neighboring organisms’ metabolism and leading to emergent interactions.

Spatio-temporal metabolic modeling begins with the development of GEMs for all constituent organisms in the simulation. GEMs are metabolic network models that include all known metabolic reactions catalyzed by enzymes encoded in the genome of a particular organism. These models can be automatically reconstructed as “draft” models based only on genome annotations (Arkin et al., 2018; Henry et al., 2010; Faria et al., 2023; Zimmermann et al., 2021; Machado et al., 2018), downloaded from curated databases (King et al., 2016; Heinken et al., 2023), or curated by hand for new organisms (Thiele and Palsson, 2010). In this chapter, we use metabolic models from the AGORA2 database, which contains semi-curated GEMs for microbial organisms commonly found in the human gut microbiome (Heinken et al., 2023).

Steady-state metabolism is simulated efficiently with flux balance analysis (FBA) (Orth et al., 2010). FBA uses linear programming to solve for the fluxes of all reactions in a given metabolic network under a given condition. FBA is typically performed by making the evolutionary assumption that microbial metabolism maximizes growth and thus maximizes the flux of the biomass reaction (a lumped pseudo-reaction that consumes biomass precursors to produce 1 gram of biomass). This maximization is subject to constraints of steady-state metabolism and reaction bounds (Equation 1). The steady-state metabolism constraint ensures that the rate of change of all metabolites is equal to zero. Thus, flux is balanced, and FBA solutions provide feasible steady-state flows through a metabolic network. The reaction bounds further constrain all fluxes to be within specified ranges. Exchange reactions are used to link the metabolic network to its environment, and the environment can be changed by updating the bounds of these exchange reactions to allow, or restrict, uptake of metabolites. In this chapter, we use a Python library for constraint-based reconstruction and analysis, COBRApy (Ebrahim et al., 2013), to load GEMs and implement FBA for the initial steady-state simulations.

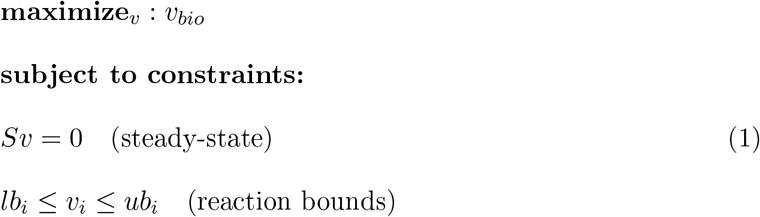

Equation 1: Steady-state simulation with flux balance analysis (FBA), where *v* is the vector of reaction fluxes, whose *i*−th component *v*_*i*_ is the flux through the reaction *i, v*_*bio*_ is the flux through the biomass reaction. Fluxes have units of mmol/(h∗g) (with g as grams dry weight of biomass), aside from the biomass reaction which produces biomass with units of g/(h∗g). The stoichiometric matrix *S* defines the stoichiometry for each reaction in the metabolic network and is used to define the steady-state constraint. Lower (*lb*) and upper (*ub*) bounds further constrain each flux.

Temporal simulation of metabolism is performed by numerically integrating the results of steady-state flux simulations. This approach is known as dynamic flux balance analysis (dFBA) (Mahadevan et al., 2002). Intracellular metabolic processes are typically assumed to be fast relative to growth and environment dynamics, allowing FBA to be applied independently at each time step. The lower bounds of the exchange reactions are constrained such that the maximum uptake of metabolites is a function of their concentration. Typically, a saturating Michaelis-Menten (or Monod) kinetic equation is used as the form of this function. Once the bounds are set, FBA is run and the abundance of biomass and metabolites in the environment are updated based on the resulting biomass and exchange fluxes (Equation 2). These simulations generate time courses of biomass and metabolite concentrations as well as time dependent metabolic fluxes (Gomez et al., 2014). In this chapter, we use the COMETS framework, without spatial context, to implement dFBA (Dukovski et al., 2021; Harcombe et al., 2014).

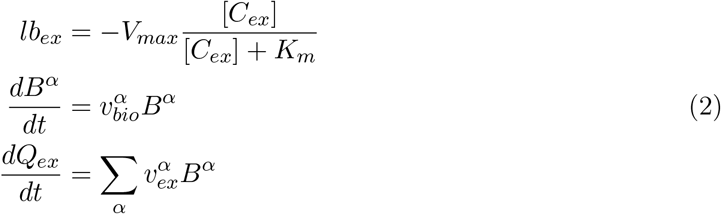

Equation 2: Temporal metabolic modeling with dynamic flux balance analysis (dFBA), where [*C*_*ex*_] is the nutrient concentration, in mmol/mL, for the exchanged metabolite *ex, V*_*max*_ is the maximum uptake rate, *K*_*m*_ is the Michaelis-Menten half-maximal uptake concentration constant, 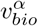 is the flux through the biomass reaction for species *α, B*^*α*^ is the biomass of species *α* in g, 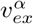 is the exchange flux for species *α* and metabolite *ex*, and *Q*_*ex*_ is the abundance of the metabolite in mmol.

Spatio-temporal simulations are implemented by extending dFBA to represent the propagation of biomass and metabolites in a structured environment using partial differential equations (Equation 3). The biomass propagation consists of two terms. The first is the diffusion of biomass. In the example we present in this chapter, we set the diffusivity of biomass as non-linear to restrict biomass diffusion to areas that are actively growing. The second term is the growth of biomass arising from the FBA simulation of steady-state metabolism. The metabolite propagation similarly consists of two terms. The first term is diffusion, which in our example is mediated by an isotropic diffusivity constant for each metabolite. The second term is the production and consumption of metabolites arising from the FBA simulation of steady-state metabolism.

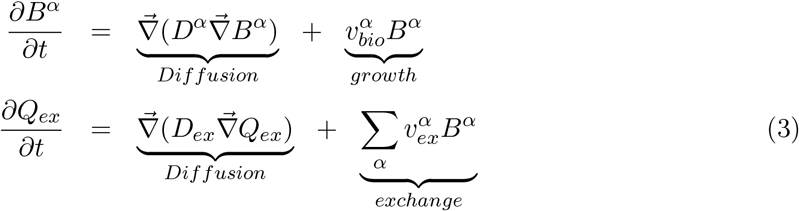

Equation 3: Spatio-Temporal metabolic modeling with partial differential equation where all variables are space (*x, y*) and time (*t*) dependent. The operator 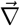 is the vector differential operator. *D*^*α*^ is the diffusivity of species *α*. In our simulations, we use a non-linear diffusivity where *D*^*α*^ = *f*(*B*^*α*^), where *f*(*B*^*α*^) is a function of the local biomass. *D*_*ex*_ is the diffusivity of external metabolites. COMETS also includes an option for advective biomass and metabolite propagation, which we do not use in this chapter. Further details on these equations and their numerical integration, as well as the form of the non-linear biomass diffusivity term, are explained in the COMETS Nature Protocols paper supplement (Dukovski et al., 2021), and subsequent work (Dukovski et al., 2025).

In this chapter, we step through a sequential process moving from steady-state, to temporal, to spatio-temporal simulation of microbial community metabolism. We provide an illustrative example that captures the interaction between bacterial colonies of *Bifidobacterium longum* subsp. *infantis* (*B. infantis*) and *Anaerobutyricum hallii* (*A. hallii*). These bacteria are native organisms of the human infant gut microbiome that have been shown to exhibit metabolic cross-feeding and contribute to short-chain fatty acid production (Schwab et al., 2017; Bunesova et al., 2018). We first identify nutrient requirements for both organisms through steady-state simulation using FBA. Next, we investigate metabolic interactions in a well-mixed environment using dFBA. Finally, we implement a spatio-temporal simulation in two spatial dimensions, using COMETS, to represent growth in the outer mucus layer of the human colon.

### 2 Materials

This section describes the computational tools that are required to implement the simulations. All code provided in this chapter is written using the Python programming language, and was developed on Mac OS 26.2 with Apple M2 chip. All analyses are also compatible with Windows and Linux operating systems given the appropriate versions of the software are installed.

### 2.1 Coding Integrated Development Environment (IDE)

All simulation code is written and executed using Jupyter notebook files (.ipynb) within Visual Studio Code. Jupyter notebook provides an interface for integrating markdown text with Python code. Visual Studio Code is an interactive computing environment that facilitates code development. While we use Visual Studio Code, all Jupyter notebook files can also be run simply using Jupyter Notebook or Jupyter Lab.

1. Visual Studio Code: https://code.visualstudio.com/Download.
2. Jupyter: https://jupyter.org/

### 2.2 Python Libraries

The simulation code uses several Python libraries (Numpy, Pandas, Matplotlib, Math, and OS) for numerical computation and system management. These libraries can be installed using pip or with a package manager such as Anaconda.

1. Numpy (Harris et al., 2020) is used for array manipulation, spatial grid operations, and general computations: https://numpy.org/.
2. Pandas (McKinney et al., 2010) is used to handle output data from COMETS simulations: https://pandas.pydata.org/.
3. Matplotlib (Hunter, 2007) is used for visualization: https://matplotlib.org/.
4. Math and OS are standard python libraries used to implement mathematical operations and manage system variables.

### 2.3 COBRApy

COBRApy is a python package that provides an interface for constraint-based reconstruction and analysis in Python (Ebrahim et al., 2013). COBRApy is used to load GEMs, analyze model properties, and run FBA.

1. COBRApy can be installed using pip or anaconda by following the instructions on the open-cobra website: https://opencobra.github.io/cobrapy/.

### 2.4 COMETS

COMETS (Computation of Microbial Ecosystem in Time and Space) is an open-source software for simulating spatio-temporal dynamics of microbial communities by integrating dFBA with biophysical models of molecular diffusion and biomass spreading (Dukovski et al., 2021; Harcombe et al., 2014). COMETS relies on a backend optimization software to implement linear programming for FBA, for which we recommend using the Gurobi optimization software.

1. COMETS can be downloaded and installed from: https://www.runcomets.org/installation In this chapter, we use version 2.12.3. Information about COMETS, including additional examples and tutorials, is available at https://www.runcomets.org.
2. Gurobi can be installed from https://www.gurobi.com/. In this chapter, we use version 10.0.3. Use of Gurobi requires a license, for which free academic licenses are currently available.

### 2.5 COMETSpy

COMETSpy is a Python library for COMETS. COMETSpy facilitates the set-up of COMETS models in Python and provides an interface to call the COMETS software to run simulations.

1. COMETSpy can be installed using pip by following the instructions in the documentation: https://cometspy.readthedocs.io/en/latest/.

### 2.6 Genome-Scale Metabolic Models (GEMs)

GEMs are computational models representing the metabolic capabilities of specific organisms. In this chapter, we use two GEMs obtained from the AGORA2 (assembly of gut organisms through reconstruction and analysis, version 2) collection (Heinken et al., 2023). We recommend using the MATLAB (.mat) version of AGORA2 GEMs which can be downloaded form the Virtual Metabolic Human website:

https://vmh.life/files/reconstructions/AGORA2/version2.01/mat_files/individual_reconstructions/.

1. Bifidobacterium_longum_infantis_ATCC_15697.mat. The reconstruction has 1032 reactions, 932 metabolites, and 163 exchange reactions. Throughout the text of this chapter, we refer to this organism as *B. infantis*.
2. Eubacterium_hallii_DSM_3353.mat (recently reclassified as *Anaerobutyricum hallii*). This model contains 1, 051 reactions, 980 metabolites, and 120 exchange reactions. Throughout the text of this chapter, we refer to this organism as *A. hallii*.

## 3 Methods

This section outlines the steps of our analysis, which builds sequentially from steady-state, to temporal, to spatio-temporal simulation of metabolism. We walk through each step and provide code excerpts. In each subsequent section, we describe only the steps/code that are substantially different from previous sections. We do not include the code used to plot figures in this chapter. The complete code is available on GitHub as Jupyter Notebook files corresponding to each section. https://github.com/m-3-lab/SpatialFBAExample/tree/main/Paper%20Code

### 3.1 Steady-State Simulation of Models and Media

The first step of the analysis is to define a representative growth medium capable of supporting the growth of all GEMs. Steady-state simulation (FBA through COBRApy) is used to identify specific metabolites that needed to be added to the medium to support growth. In our example, we used this step to identify a minimal glucose medium that supports the growth of *B. infantis* and *A. hallii* while encouraging metabolic interactions.

1. Import the libraries. Import COBRApy to run FBA. 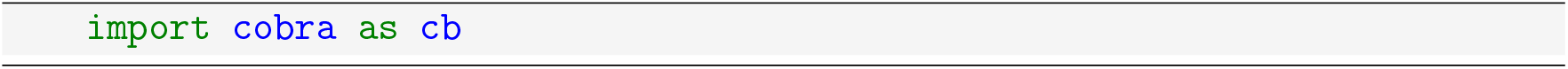
2. Load the models. Download the GEMs for *B. infantis* and *A. hallii* from the AGORA2 database, as described in the materials, and place the model (.mat) files in the local directory. Use COBRApy to import the model files into Python objects (*see* **Note 1**). 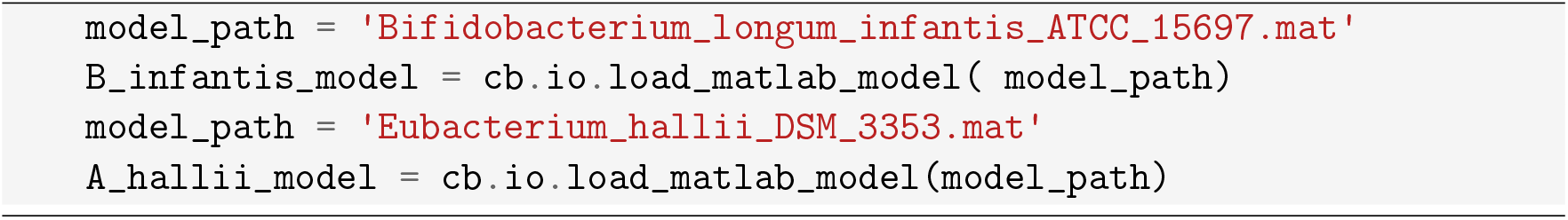
3. Define the base medium. This medium contains metabolites that are known to be present in the environment. The medium is defined as a dictionary that maps exchange reaction IDs to maximum uptake rates (absolute value of the exchange lower bounds in mmol/(g∗h)) (*see* **Note 2**). 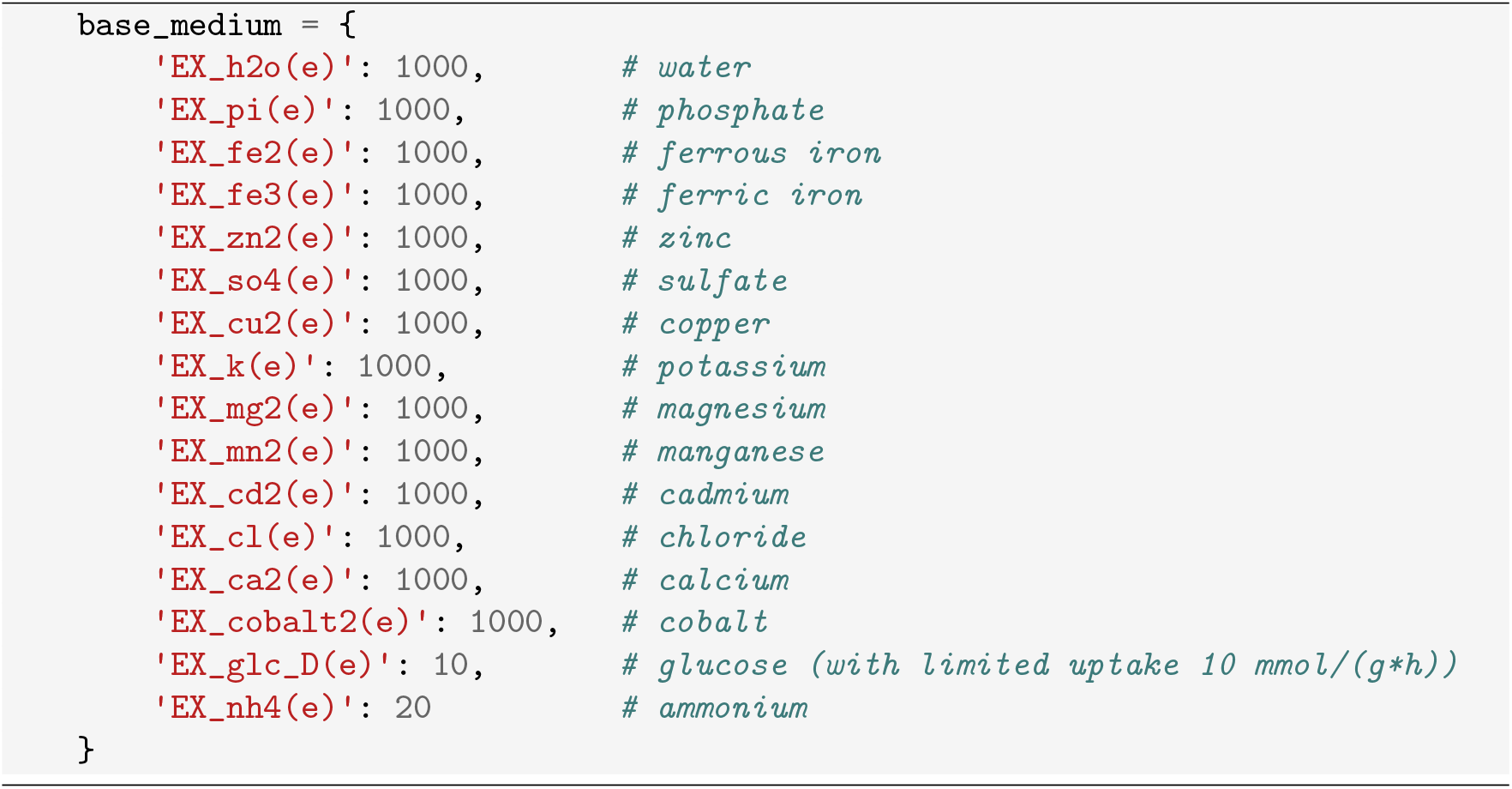
4. Find the minimal medium for (*B. infantis*). We developed a modified implementation of the COBRApy “minimal_medium” function, which allows us to first provide the base medium (using temporary sink reactions), and then identify the minimal number of metabolites that need to be added to the base medium to enable growth of the model. The output of this code provides a list of metabolites that were added to the medium to enable growth. The “minimal_medium” function identifies “minimize_components” number of suggested new medium compositions. To define a minimal medium, iteratively run the modified minimal medium code, examine the output, and add new components to the medium. In our example, we added a small number of suggested metabolites to complete the medium while avoiding the addition of oxygen to ensure an anaerobic environment. 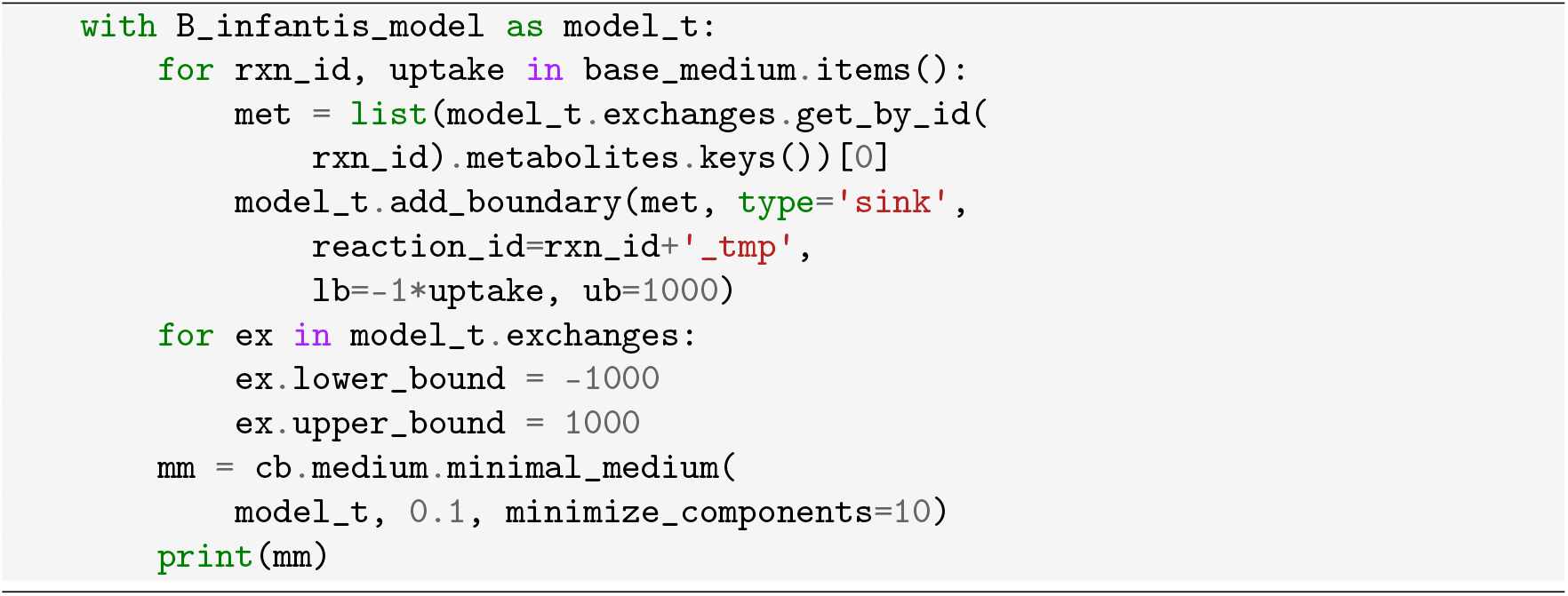
5. Define the new growth medium for *B. infantis*. From iterative application of the above step, identify a minimal growth medium that supports the growth of (*B. infantis*) and define a new medium. 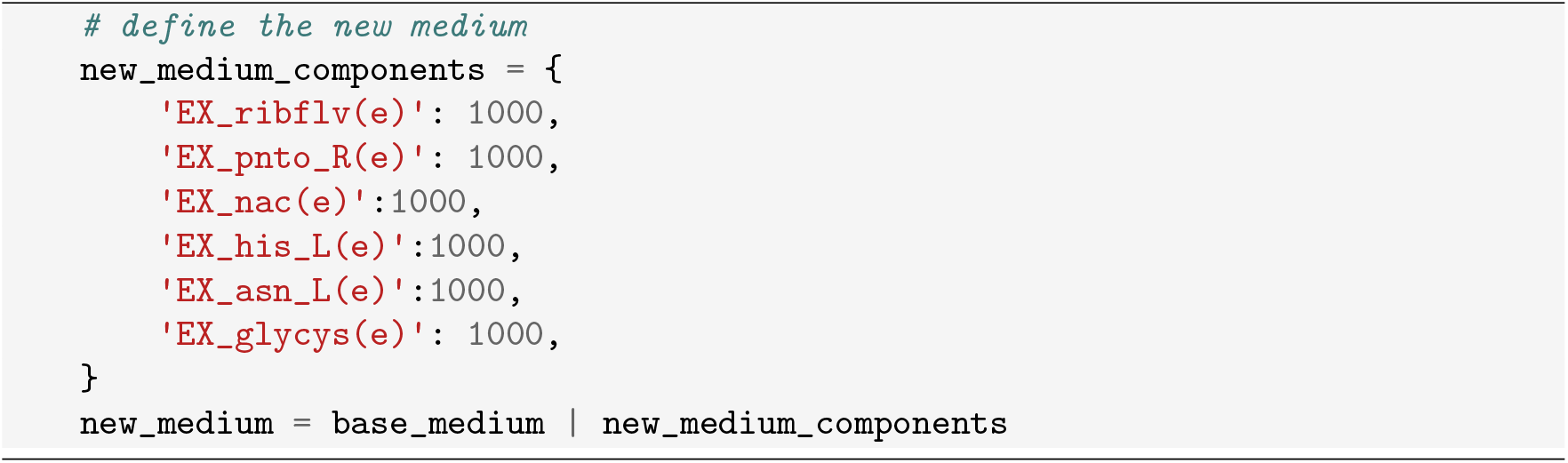
6. Confirm *B. infantis* growth on the new medium. Run FBA to confirm *B. infantis* growth (*sol*.*objective*_*value*) is positive. The expected growth rate is 0.11 h^−1^. Print the value of the L-lactate exchange flux (*EX*_*lac*_*L*(*e*)). The expected L-lactate production is 18.4 mmol/(g∗h). In our example, we anticipated that cross-feeding would be driven by L-lactate and thus wanted to confirm L-lactate production. 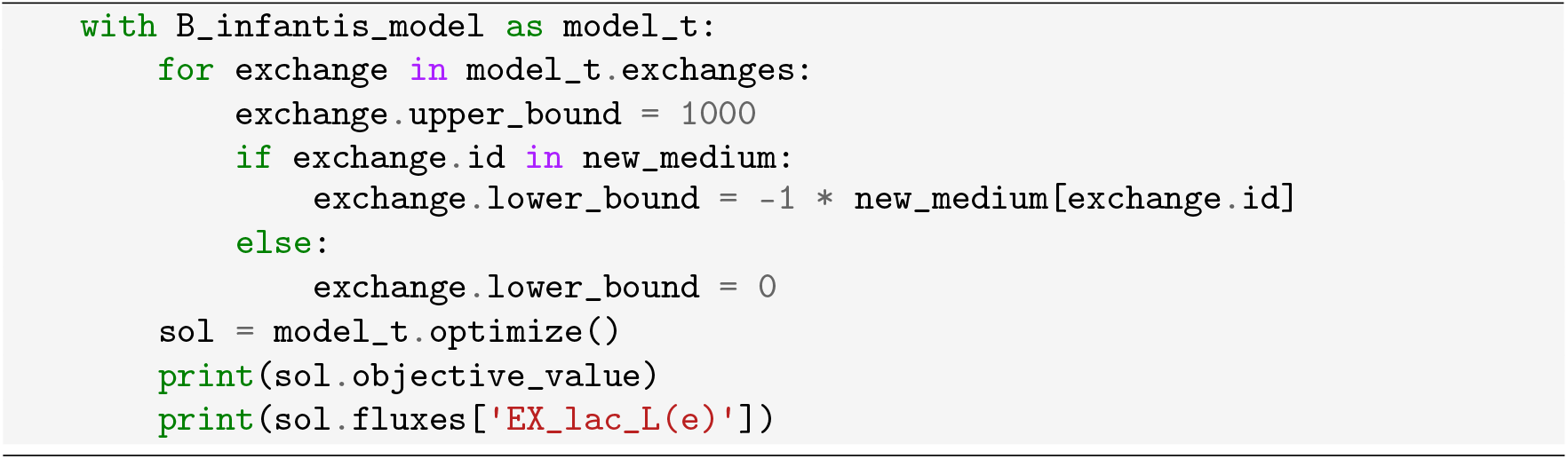
7. Identify additional medium components for *A. hallii*. Add L-lactate to the medium to encourage cross-feeding with *B. infantis*. Re-run the modified “minimal_medium” function with the new base medium to identify additional components that are needed for *A. hallii* growth. 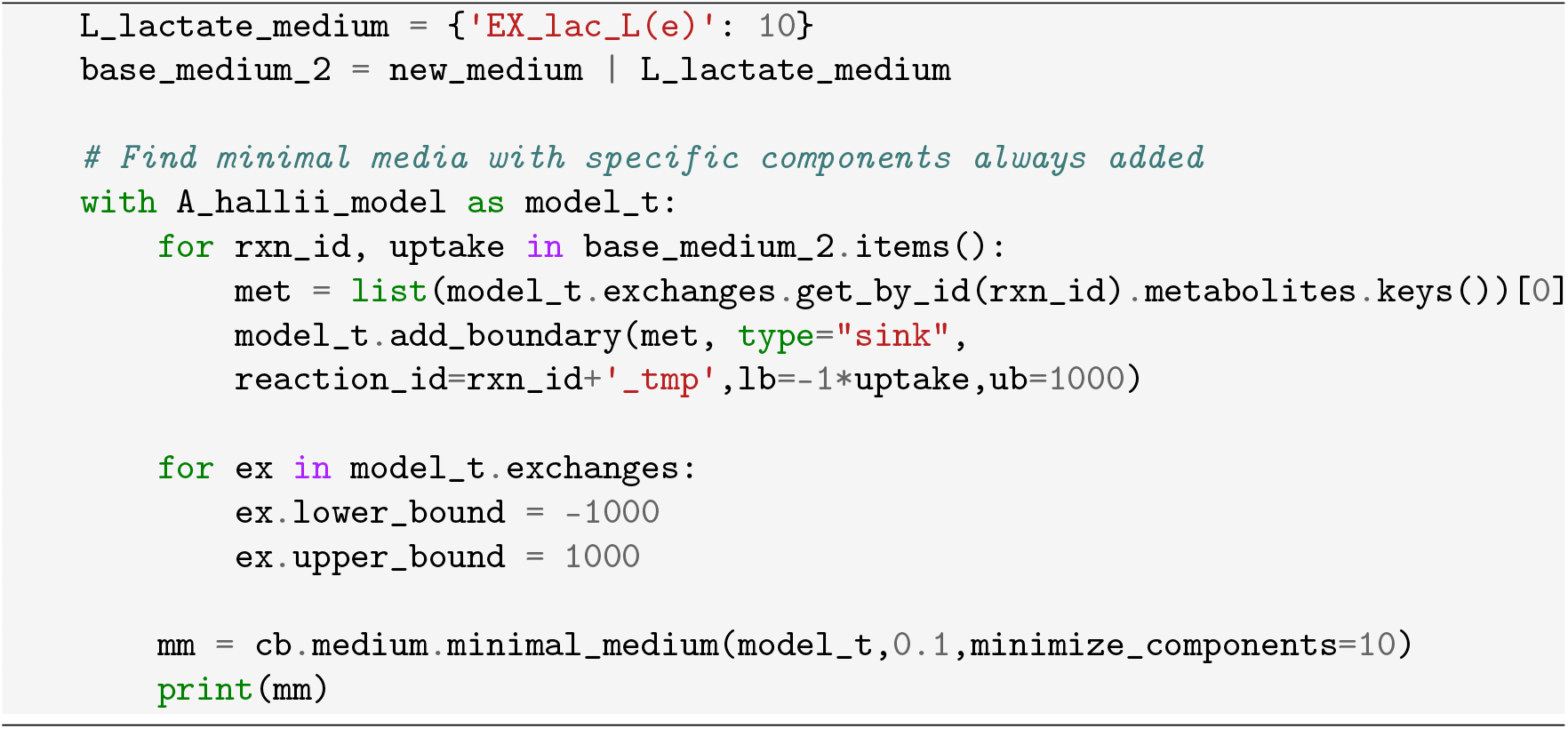
8. Define the final medium, enabling both *B. infantis* and *A. hallii* anaerobic growth. 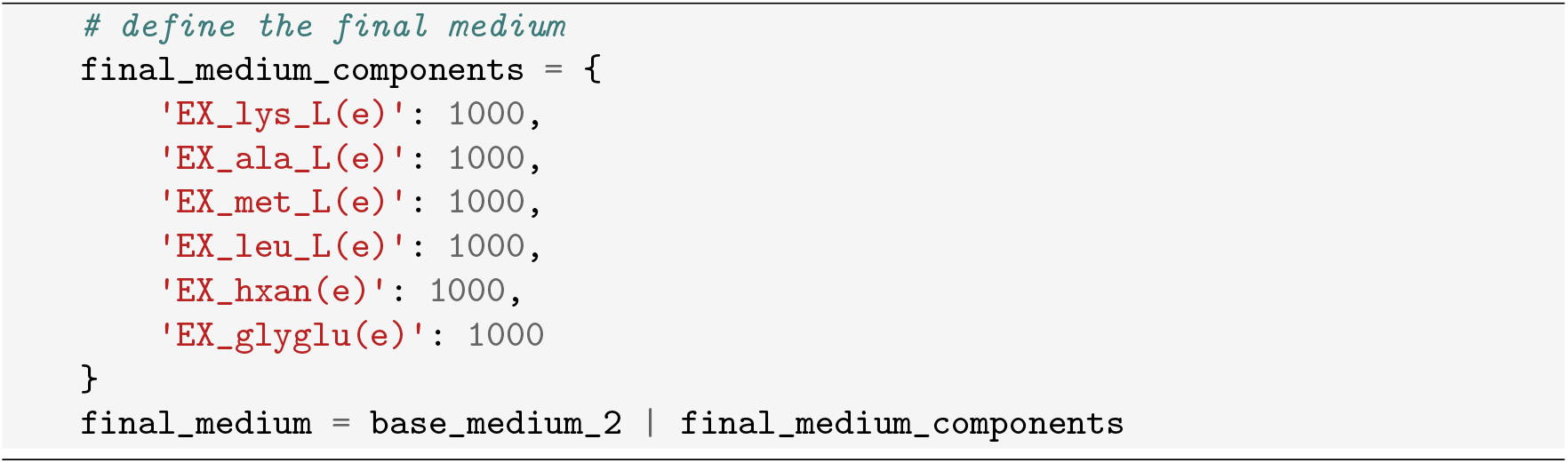
9. Confirm growth of both organisms in the final medium with FBA, as done in step 6. The expected growth rate for *B. infantis* is 0.13 h^−1^ and for *A. hallii* is 0.50 h^−1^.

**Table 1:** Final medium used for steady-state flux simulations. Uptake bounds are used to set the negative value of the lower bound of the corresponding exchange reaction, with units of mmol/(g*h). The medium consists of metabolites added from the base medium, those added to ensure *B. infantis* growth, those added to ensure *A. hallii* growth, and L-lactate to examine cross-feeding.

| Exchange_Reaction | Metabolite_ID | Metabolite_Name | Uptake_Bound | Notes |
| --- | --- | --- | --- | --- |
| EX_h2o(e) | h2o[e] | Water | 1000 | Base |
| EX_pi(e) | pi[e] | hydrogenphosphate | 1000 | Base |
| EX_fe2(e) | fe2[e] | Fe2+ | 1000 | Base |
| EX_fe3(e) | fe3[e] | Fe3+ | 1000 | Base |
| EX_zn2(e) | zn2[e] | Zinc | 1000 | Base |
| EX_so4(e) | so4[e] | sulfate | 1000 | Base |
| EX_cu2(e) | cu2[e] | Cu2+ | 1000 | Base |
| EX_k(e) | k[e] | potassium | 1000 | Base |
| EX_mg2(e) | mg2[e] | magnesium | 1000 | Base |
| EX_mn2(e) | mn2[e] | Mn2+ | 1000 | Base |
| EX_cd2(e) | cd2[e] | Cadmium | 1000 | Base |
| EX_cl(e) | cl[e] | Chloride | 1000 | Base |
| EX_ca2(e) | ca2[e] | calcium(2+) | 1000 | Base |
| EX_cobalt2(e) | cobalt2[e] | Co2+ | 1000 | Base |
| EX_glc_D(e) | glc_D[e] | D-glucose | 10 | Base |
| EX_nh4(e) | nh4[e] | Ammonium | 20 | Base |
| EX_ribflv(e) | ribflv[e] | Riboflavin | 1000 | B. infantis |
| EX_pnto_R(e) | pnto_R[e] | (R)-Pantothenate | 1000 | B. infantis |
| EX_nac(e) | nac[e] | Nicotinate | 1000 | B. infantis |
| EX_his_L(e) | his_L[e] | L-histidine | 1000 | B. infantis |
| EX_asn_L(e) | asn_L[e] | L-asparagine | 1000 | B. infantis |
| EX_glycys(e) | glycys[e] | Gly-Cys | 1000 | B. infantis |
| EX_lac_L(e) | lac_L[e] | L-lactate | 10 | Secreted by B. infantis |
| EX_lys_L(e) | lys_L[e] | L-lysine | 1000 | A. hallii |
| EX_ala_L(e) | ala_L[e] | L-alanine | 1000 | A. hallii |
| EX_met_L(e) | met_L[e] | L-methionine | 1000 | A. hallii |
| EX_leu_L(e) | leu_L[e] | L-leucine | 1000 | A. hallii |
| EX_hxan(e) | hxan[e] | Hypoxanthine | 1000 | A. hallii |
| EX_glyglu(e) | glyglu[e] | Glycyl-L-glutamate | 1000 | A. hallii |

### 3.2 Temporal Simulation in a Well-Mixed Environment

This section provides the steps to setup and simulate the individual (monoculture) and community (co-culture) temporal dynamic flux balance analysis (dFBA) simulations using COMETS. The minimal glucose medium identified in step 3.1 is used to define the components of the environment with the concentrations of metabolites set to physiologically relevant levels. In our example, we simulated *B. infantis* and *A. hallii* mono-cultures (3.2.1) and co-cultures (3.2.2), and further investigated the dynamics of *A. hallii* butyrate production across varying amounts of input L-lactate and glucose (3.2.3).

#### 3.2.1 Individual Simulations

This section outlines the process of simulating temporal dynamics for mono-cultures of *B. infantis* and *A. hallii*.

1. Configure COMETS and Gubrobi system paths at the top of the notebook (*see* **Note 3**). 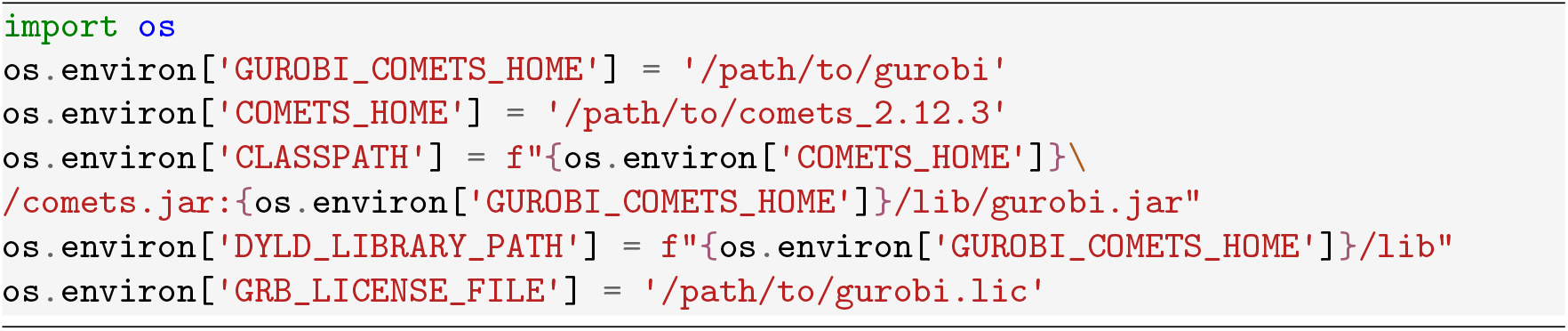
2. Import Python libraries. 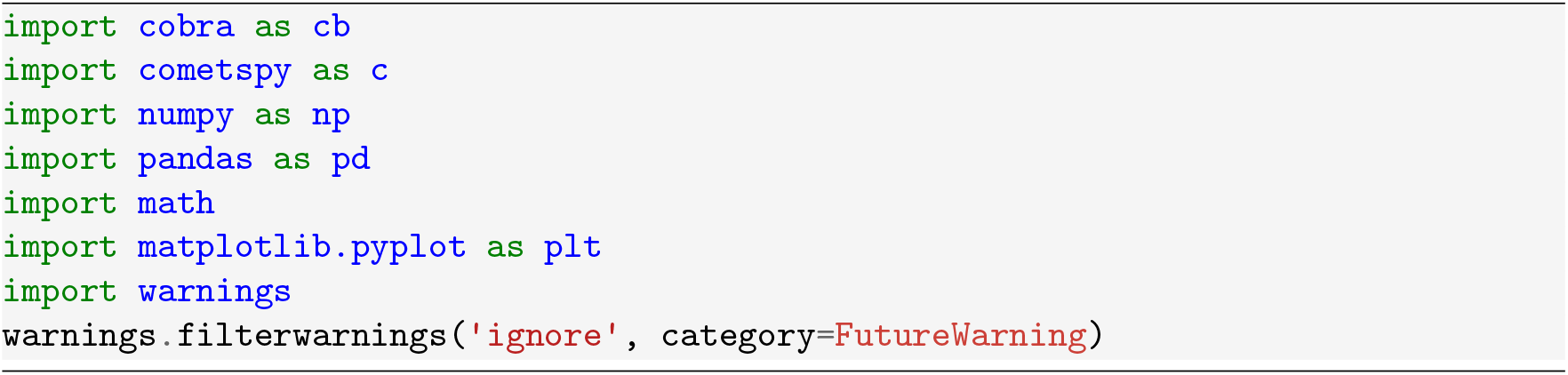
3. Define the simulation geometry. The well-mixed temporal simulations use a single 1 × 1 grid cell, and this step defines the geometry of that cell (*see* **Note 4**). 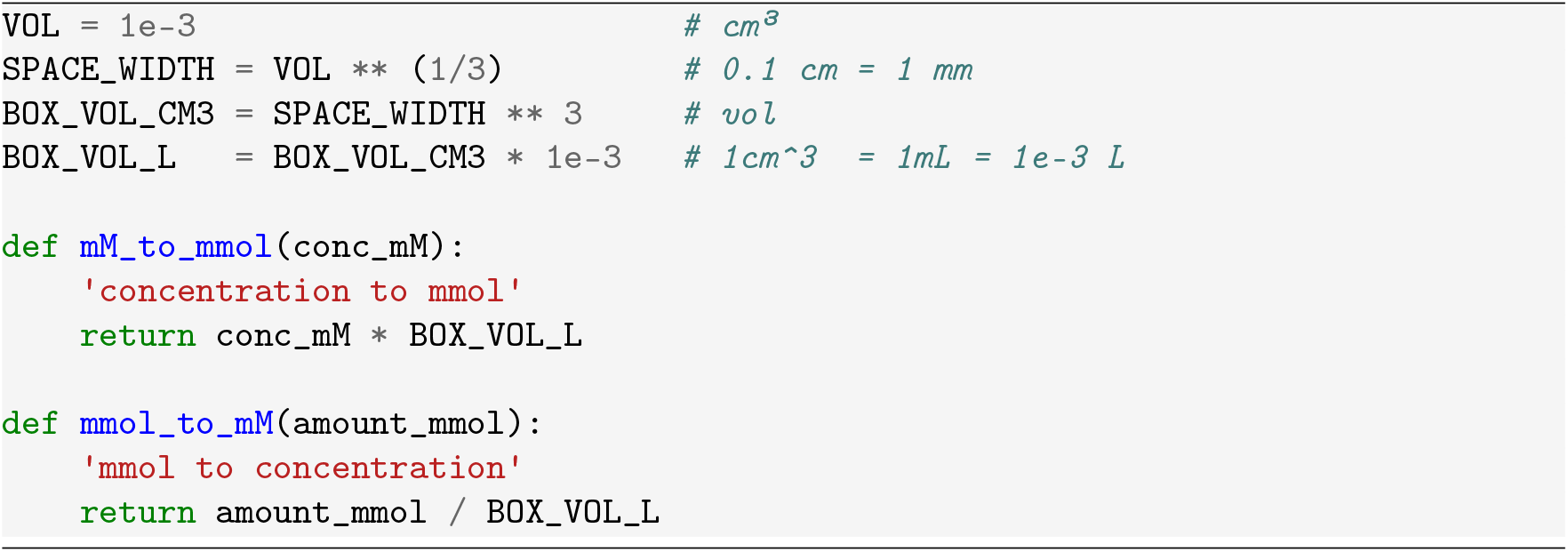
4. Define the COMETS simulation parameters (*see* **Note 5**). Define concentrations or amounts for major classes of medium metabolites. 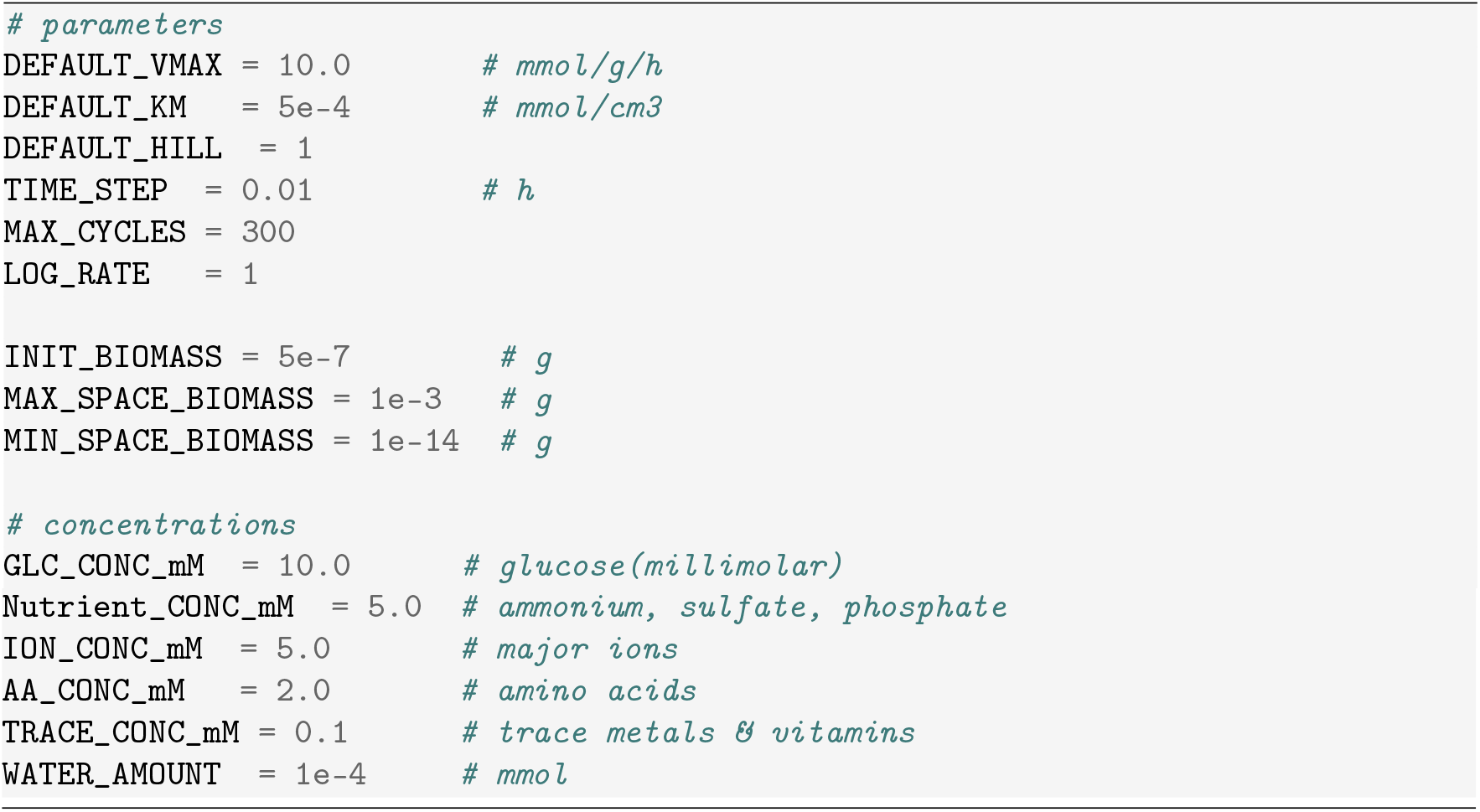
5. Define combined medium from section 3.1. The medium is assumed to be glucose limited. Glucose is set as a depletable metabolite in the simulation. All other medium components, defined here, are set at as static metabolites. Static metabolites are always provided at a defined concentration. The medium here is organized into different categories of metabolites, which can be provided at different static concentrations. 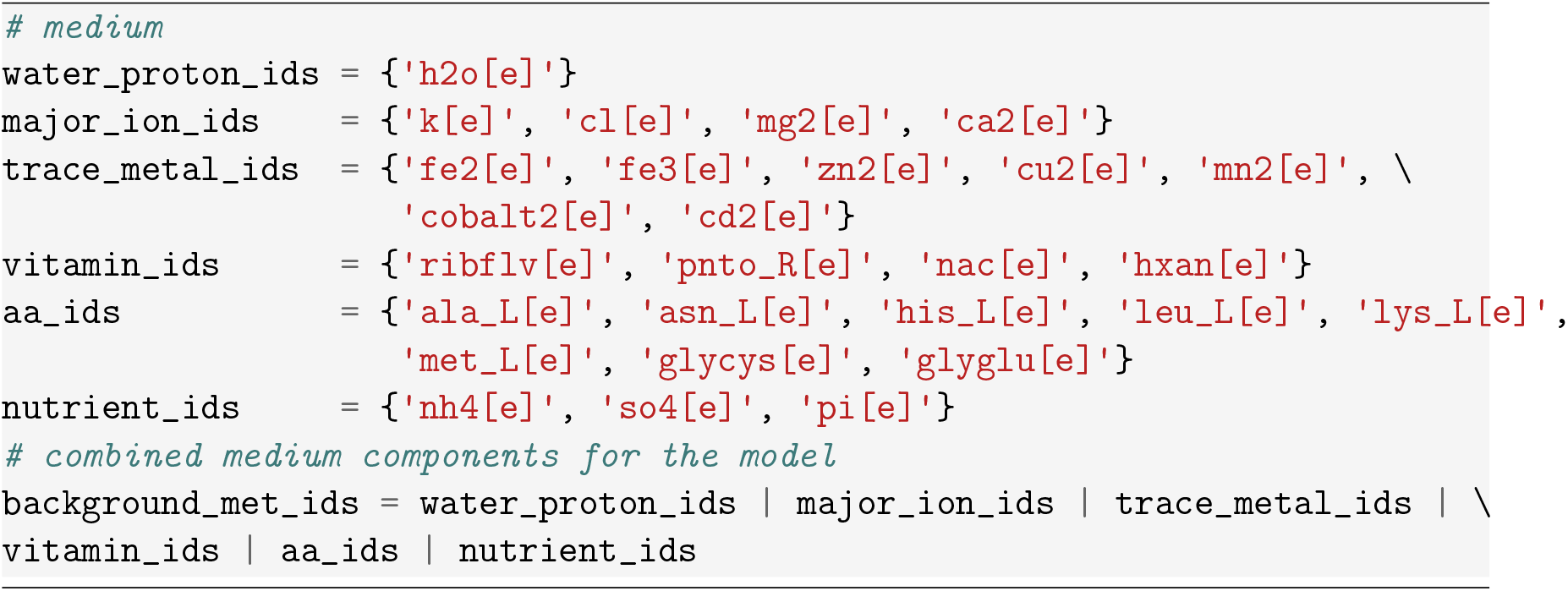
6. Load *B. infantis* model, create a COMETS object, open all exchanges and set initial biomass. Open exchanges allows the COMETS model to uptake any metabolite that is produced during the simulation. Clean non-exchanges with the “clean_non_ex” function (*see* **Note 6**). 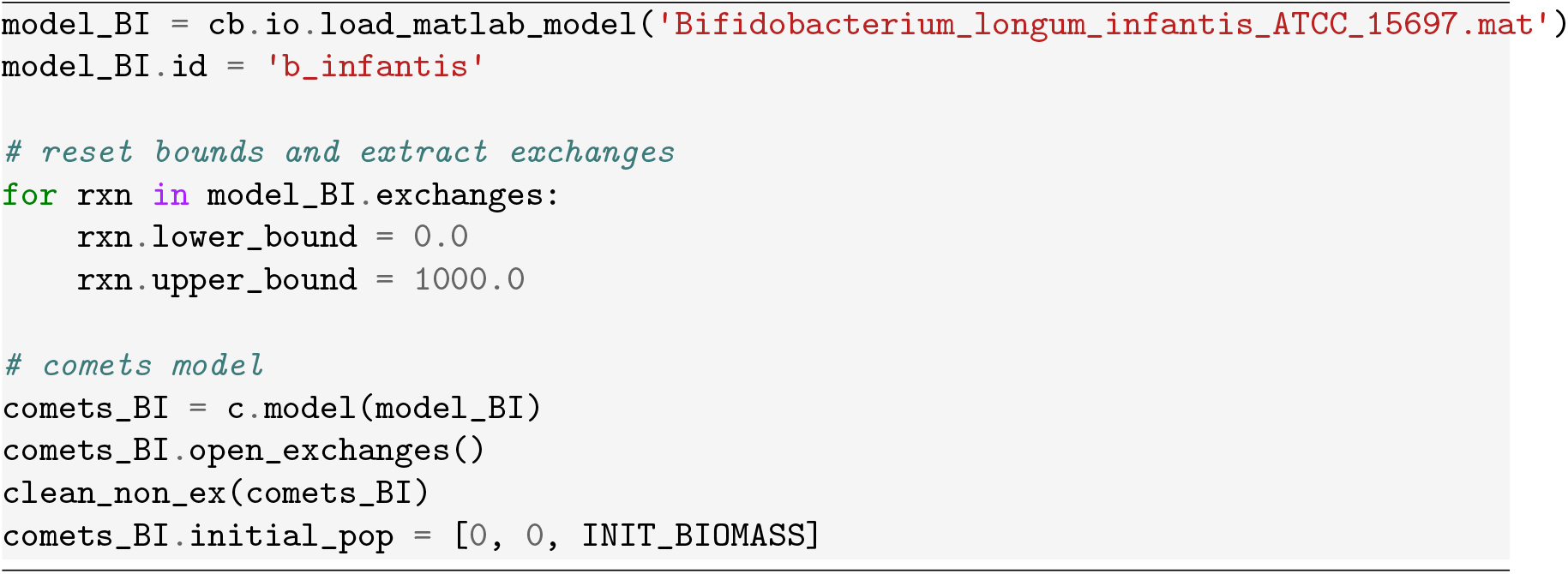
7. Create the layout. Set the medium at static concentrations using the “classify_and_set_media” function. This function sets the concentration of all exchanged metabolites in the model (“all_ex_mets_BI”) to 0 unless they are specified in the medium. Configure glucose as a depletable metabolite. 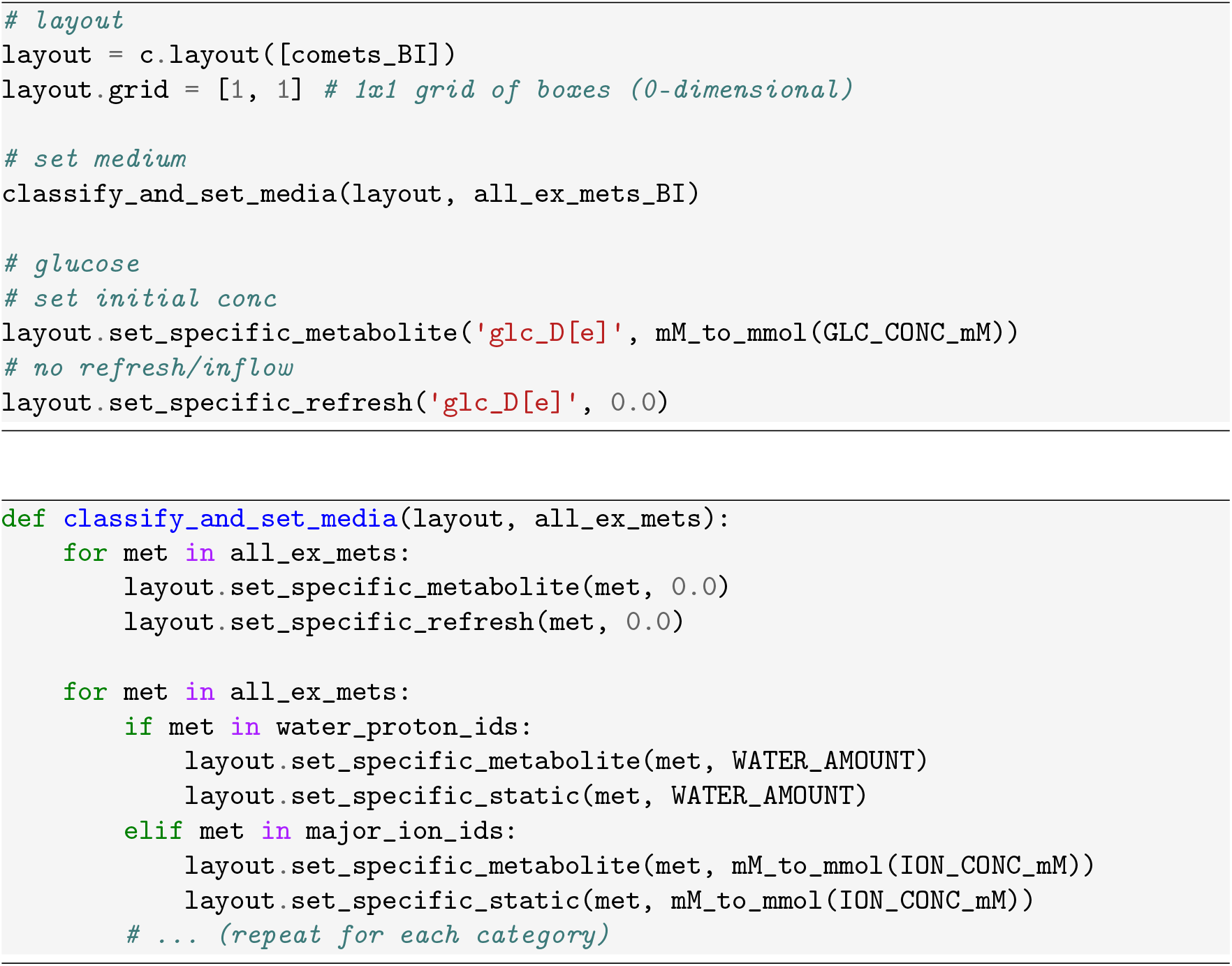
8. Build the parameter set, combine the layout and parameters into a COMETS object, and run the simulation. These mono-culture simulations take around 5 seconds to run. 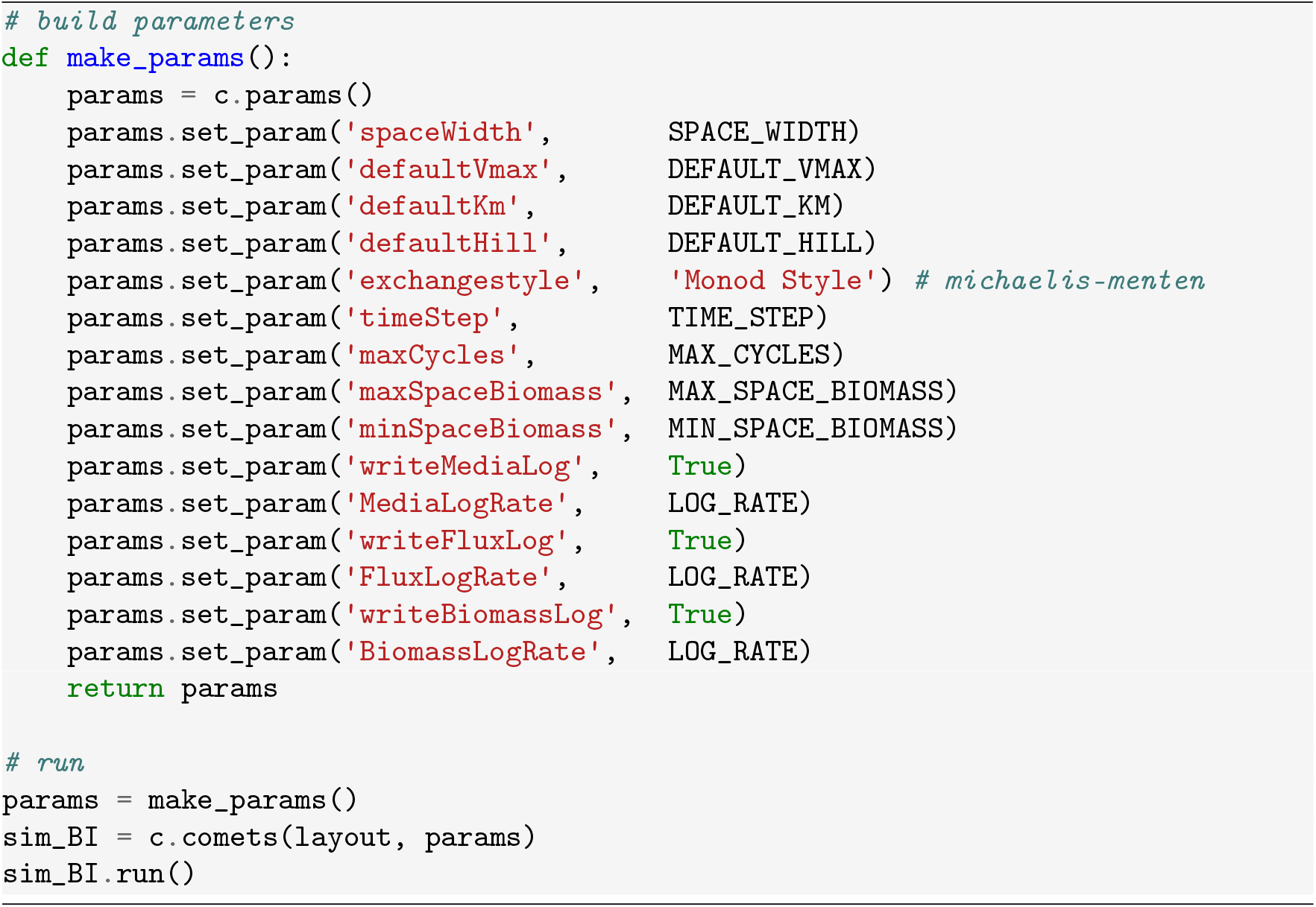
9. Extract results from the completed simulation objects and plot simulation dynamics (Figure 1). 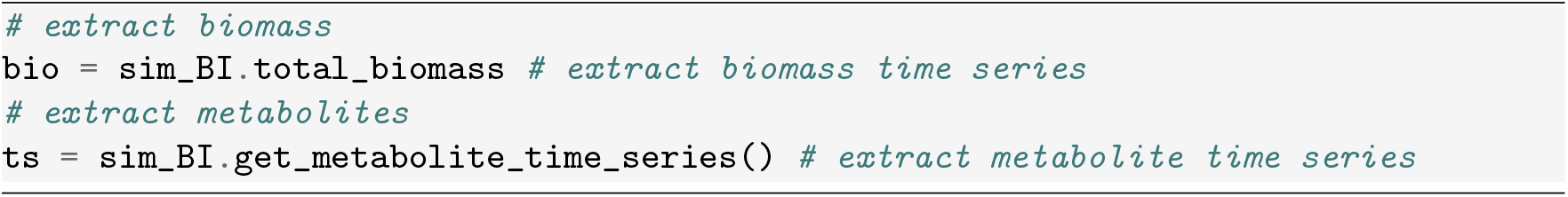
10. Repeat the same process for *A. hallii* mono-culture, loading the *A. hallii* model, running under identical conditions, and extracting and plotting results (Figure 2).

**Figure 1:**
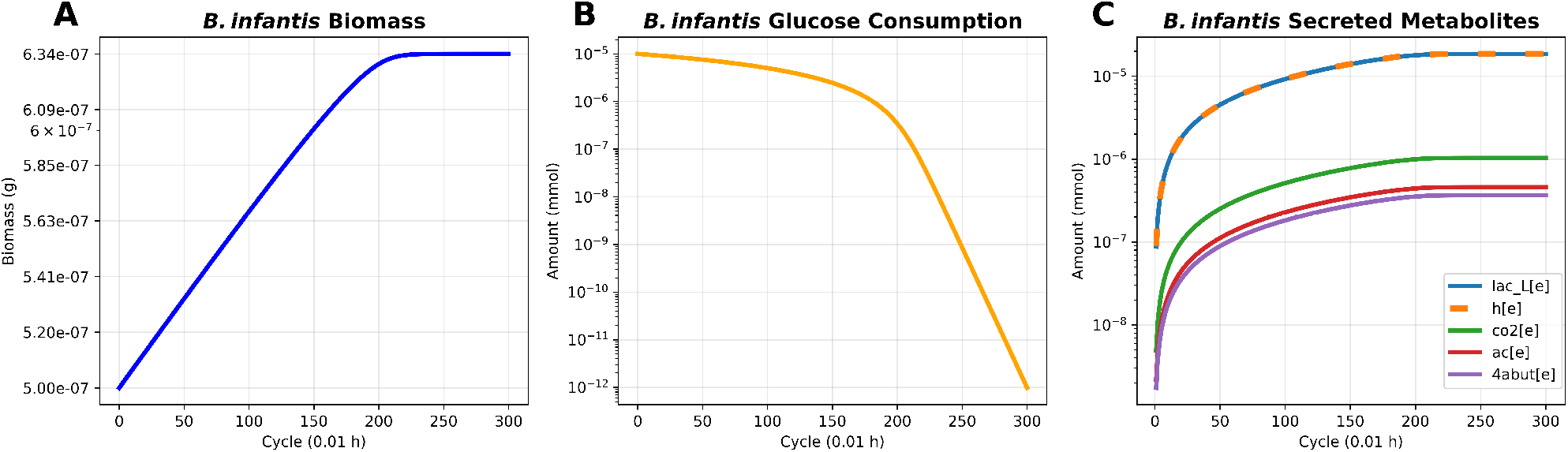
*B. infantis* monoculture dynamics. (A) Biomass growth. *B. infantis* shows exponential growth on glucose, with the biomass increasing from an initial amount of 5 × 10^−7^ g. (B) Glucose consumption. Glucose is consumed progressively over the simulation period and limits growth. (C) Secreted metabolites. The metabolite secretion profile shows L-lactate as the dominant product of interest. All plots use a log y-axis.

**Figure 2:**
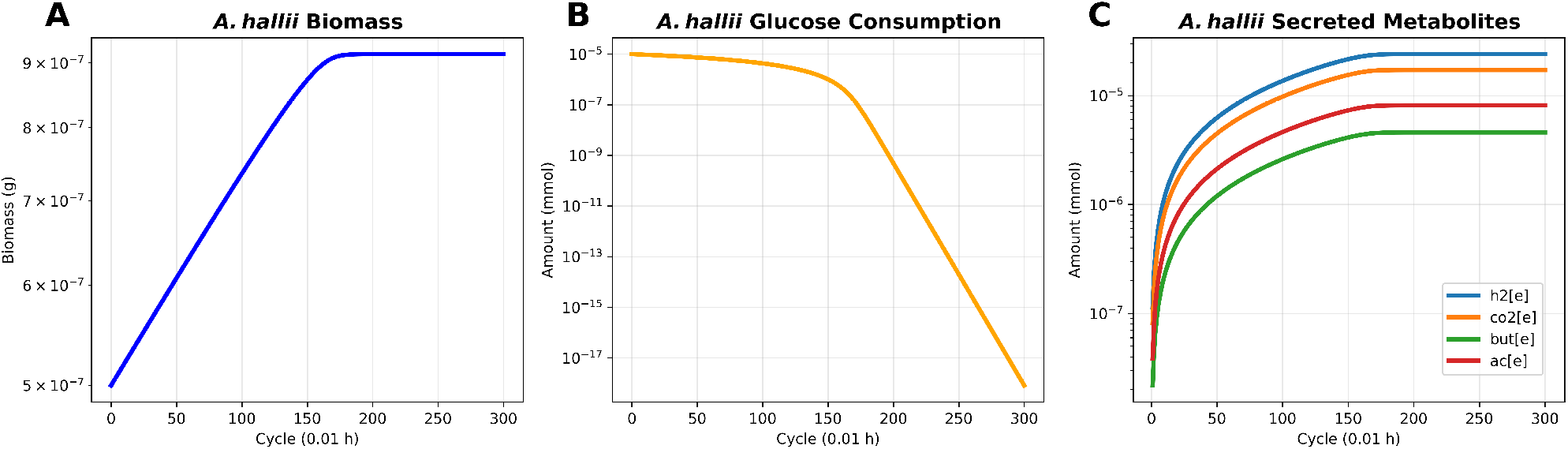
*A. hallii* monoculture dynamics. (A) Biomass growth. *A. hallii* shows exponential growth on glucose, with the biomass increasing from an initial amount of 5 × 10^−7^ g. (B) Glucose consumption. Glucose is consumed progressively over the simulation period and limits growth. (C) Secreted metabolites. The short-chain fatty acids butyrate and acetate are produced. All plots use a log y-axis

#### 3.2.2 Co-culture Simulation

This section outlines the process of running a two-species co-culture simulation in a well-mixed environment. The goal of our simulation is to observe emergent competition and cross-feeding behavior between *B. infantis* and *A. hallii*. Mono-culture simulations of each organism are also re-run for comparison.

1. Modify the simulation parameters in the monoculture for the co-culture. Reduce the simulation to 150 cycles, increase the maximum biomass per box to 10^−2^ g, and increase the maximum uptake rate to 20 mmol/(g∗h) to efficiently simulate strong interactions. 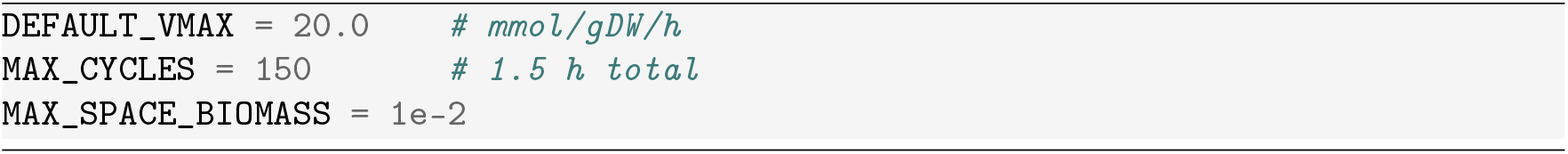
2. Load and modify GEMs. Ensure that the GEMs have matching exchange reactions by adding closed exchanges (*see* **Note 7**). Create COMETS models and open exchanges. Place both species in the same box (0,0) with biomass 5 × 10^−7^ g each to simulate a well mixed environment. 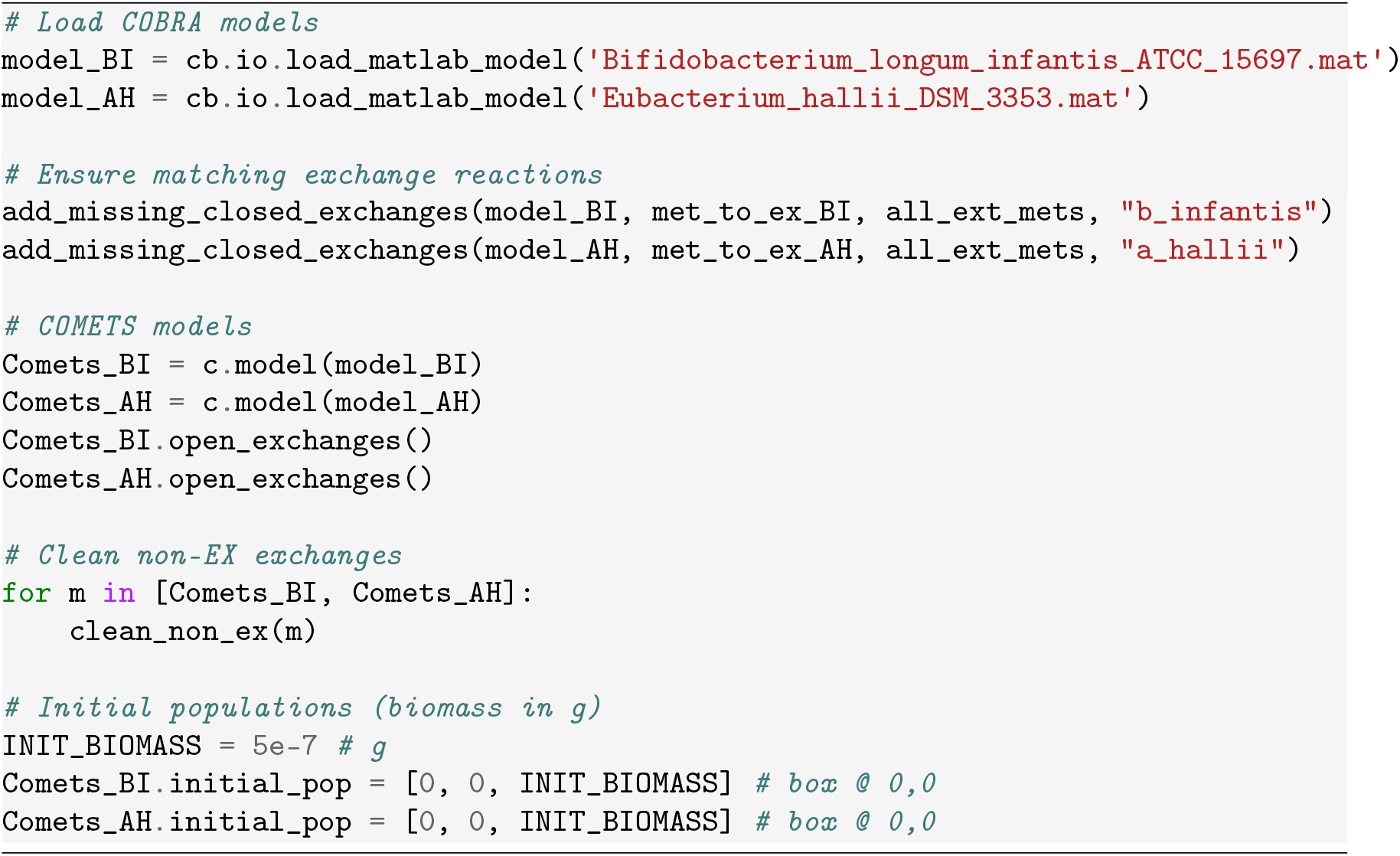
3. Create a unified layout function that can handle both monoculture and co-culture simulations. 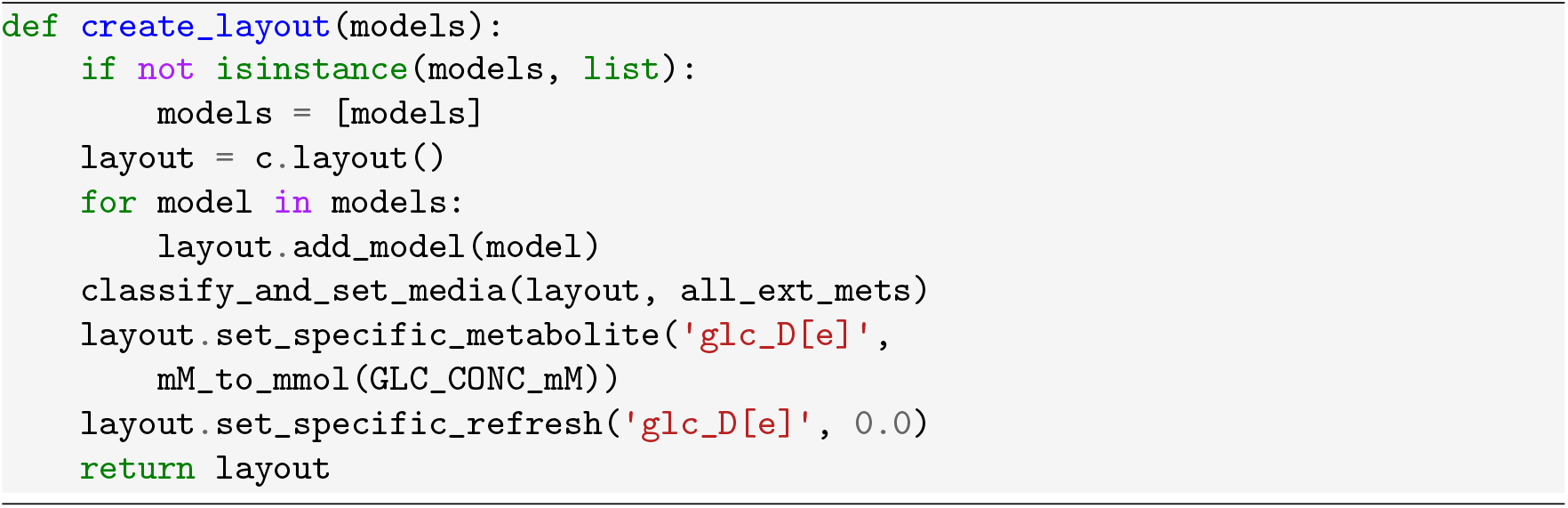
4. Run three simulations: *B. infantis* monoculture, *A. hallii* monoculture, and co-culture with both species. These simulations combined take around 11 seconds to run. 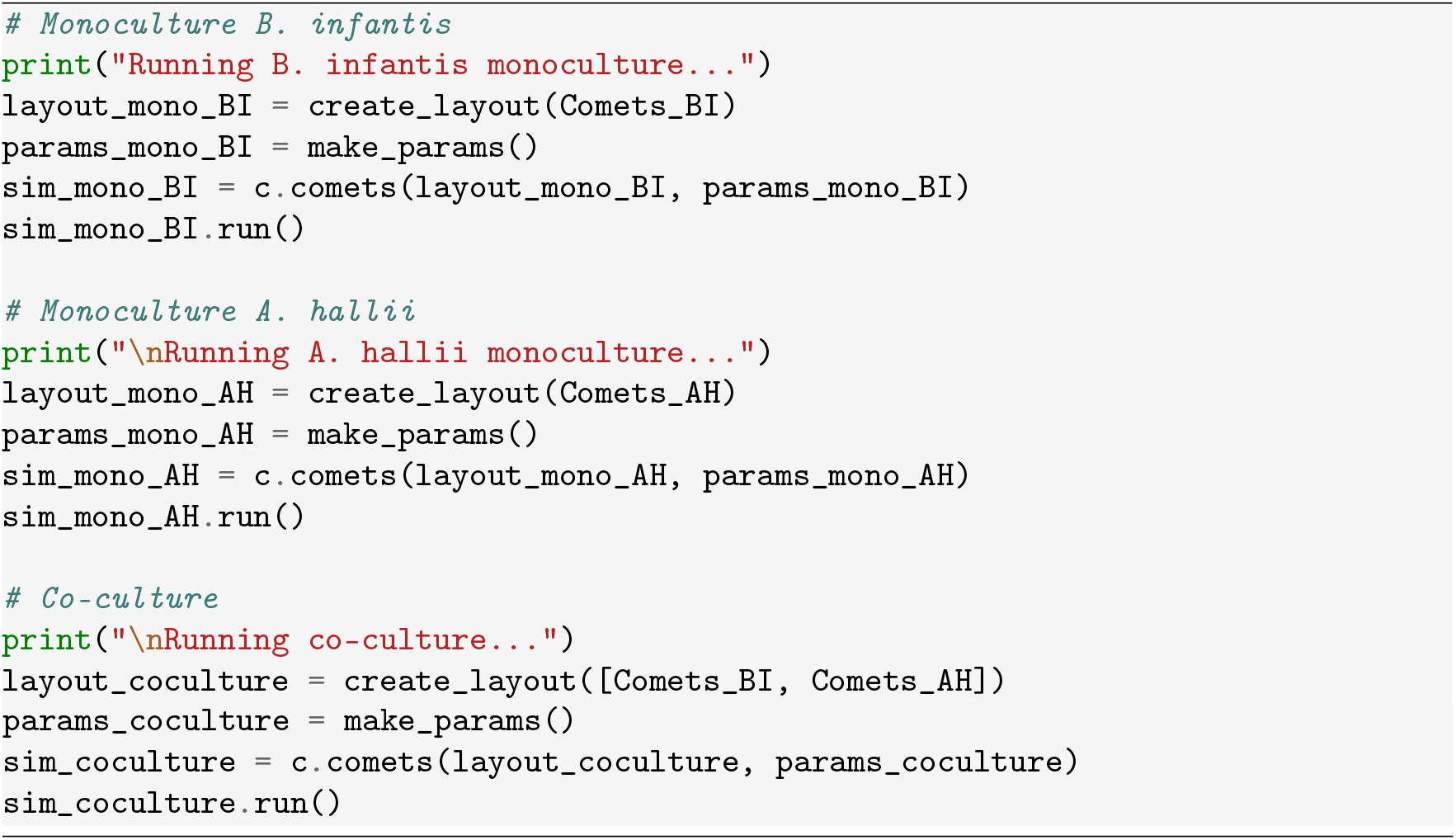
5. Compare biomass and metabolite dynamics from mono-cultures and co-culture (Figure 3).
6. Analyze the exchange fluxes. To gain further insight into the metabolic dynamics, extract the flux data using “sim.get_species_exchange_fluxes()” (Figure 4).

**Figure 3:**
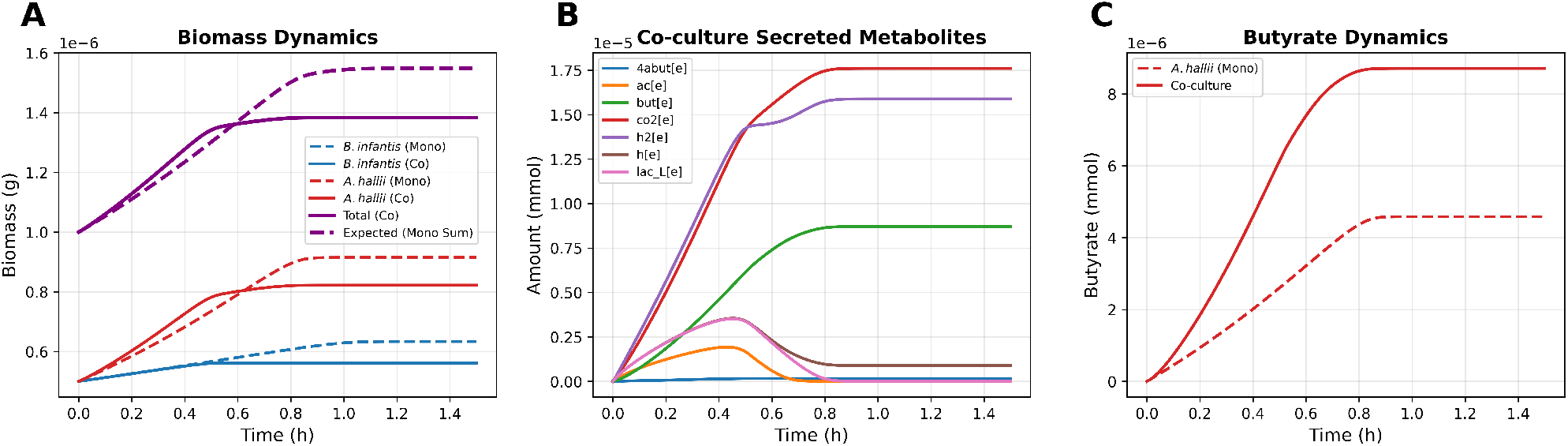
Co-culture simulation results. (A) Biomass dynamics compared between mono-culture and co-culture for both species. *A. halli* grows to a higher biomass than *B. infantis* in both mono-culture and co-culture. Both organisms grow to a lower yield in co-culture relative to mono-culture, and the final total biomass in co-culture is less than the sum of individual biomass in mono-culture. This indicates that competition limits overall growth. (B) Secreted metabolite profiles in co-culture. L-lactate and acetate peak and later decline, suggesting cross-feeding. (C) Butyrate dynamics compared between *A. hallii* mono-culture and co-culture. Butyrate production is higher in co-culture, compared to mono-culture, despite the decrease in *A. hallii* biomass.

**Figure 4:**
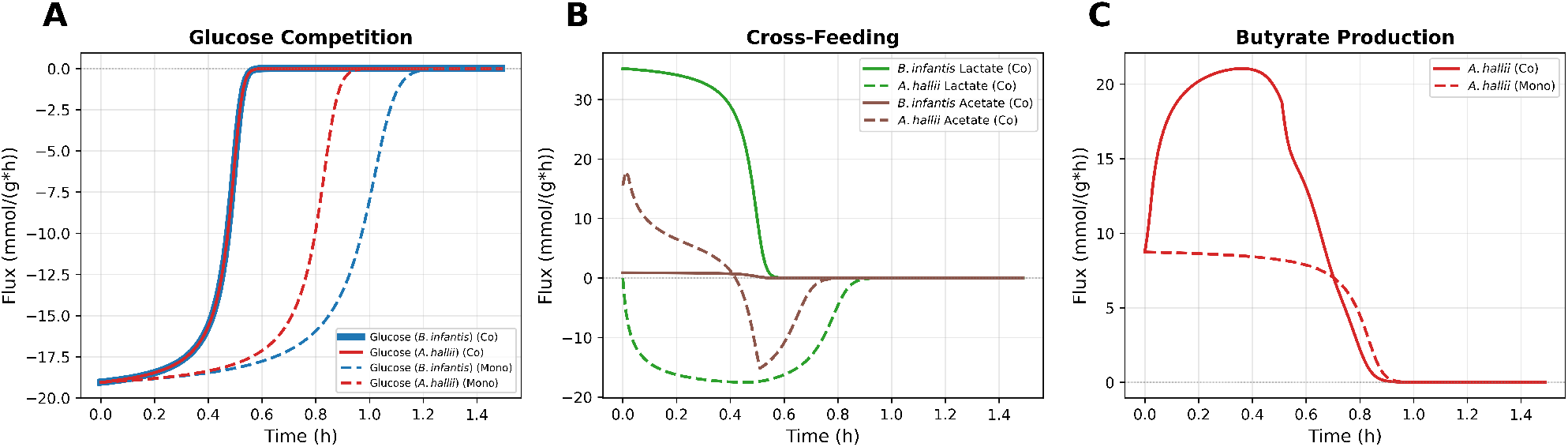
Co-culture exchange fluxes. (A) Uptake of glucose in co- and mono-cultures. Glucose is consumed by both organisms (negative exchange flux), and simultaneous uptake of this metabolite leads to competition that limits growth. (B) L-lactate and acetate exchange fluxes in co-culture. L-lactate fluxes demonstrate strong cross-feeding from *B. infantis* (producing) to *A. hallii* (consuming). Acetate is produced at low levels by *B. infantis* and is produced at early time points and consumed at later time points by *A. hallii*. (C) Butyrate production by *A. hallii* in co- and mono-cultures. Butyrate production flux is higher in co-culture relative to the *A. hallii* mono-culture. This increase corresponds with *A. hallii* uptake of L-lactate in co-culture.

#### 3.2.3 Butyrate Production Analysis

The co-culture results suggested that both glucose and L-lactate consumption contribute to butyrate production by *A. hallii*. We were interested in further understanding these dynamics by running additional simulations of *A. hallii* metabolism in defined media with varying amounts of L-lactate and glucose. This section describes a systematic parameter sweep to determine the optimal ratio of L-lactate to glucose that maximizes butyrate production while maintaining the total amount of carbon input into the system.

1. Define the parameter sweep range. Generate 21 conditions by varying glucose from 10 mM to 2 mM and the L-lactate from 0 to 16 mM. Glucose provides 6 carbons per molecule and L-lactate provides 3 carbon per molecule. The total carbon provided is kept constant at 60 mM for all parameter values. 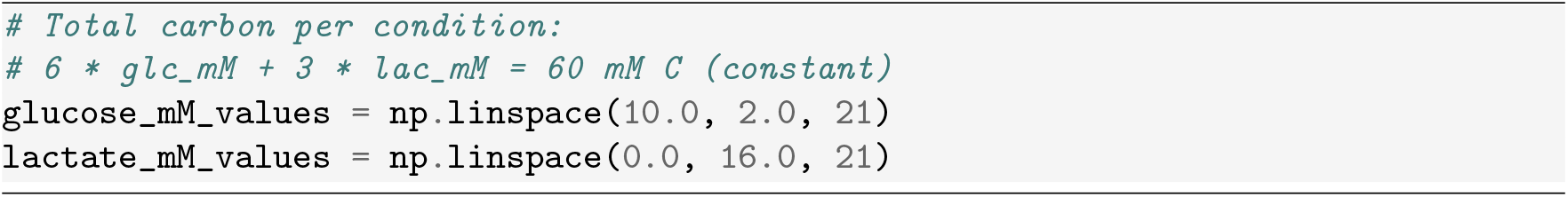
2. Define a function that sets up and runs a single simulation for a given glucose and L-lactate concentration. 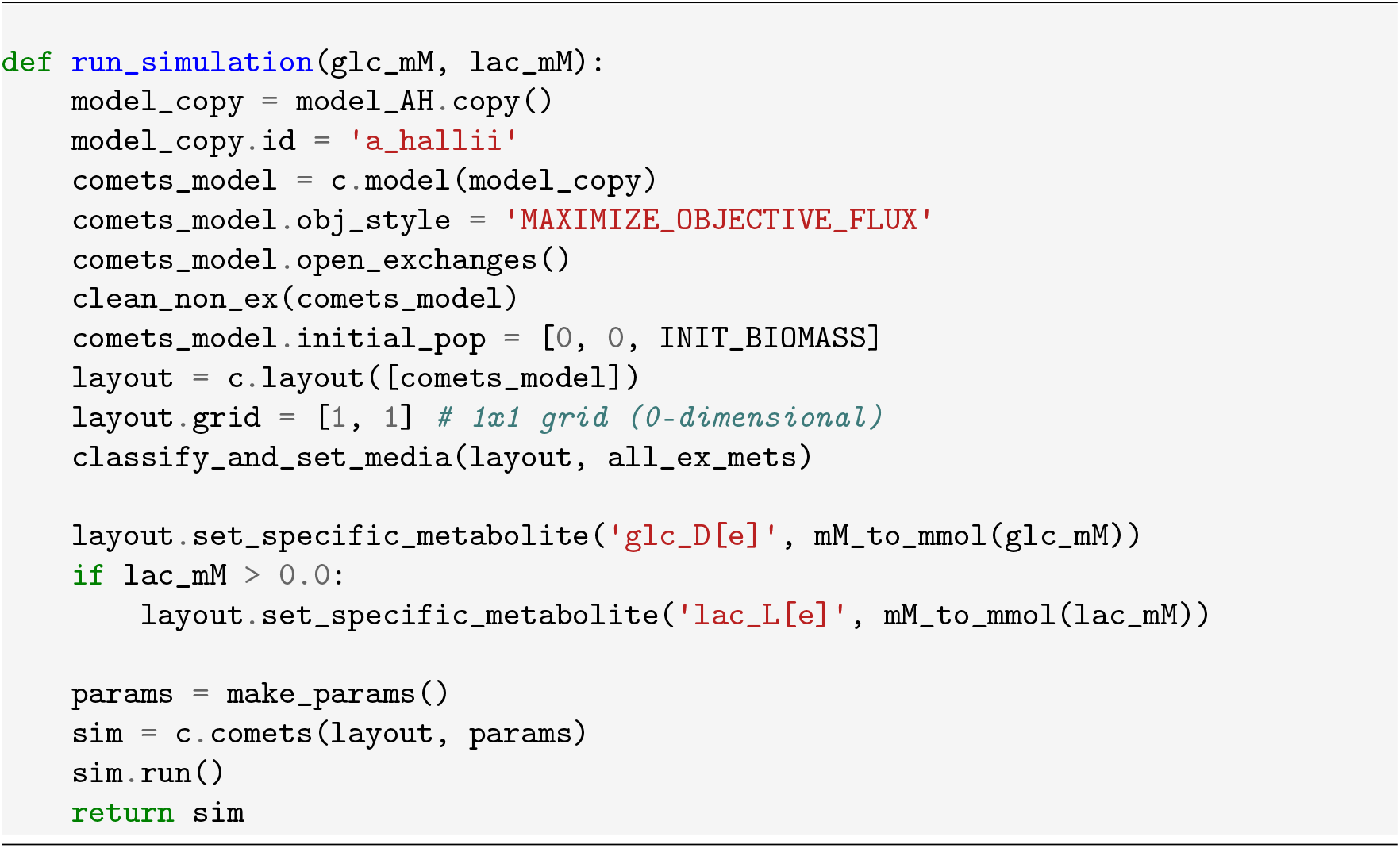
3. Run all 21 conditions sequentially. These simulations take around 96 seconds to run. 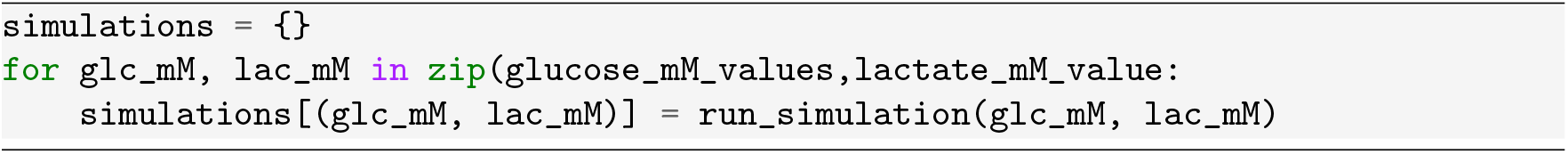
4. Analyze *A. hallii* biomass and butyrate production for each condition. Plot biomass and butyrate production as functions of the L-lactate carbon percentage (Figure 5). The L-lactate carbon percentage is calculated from the initial L-lactate and glucose concentration in each condition: 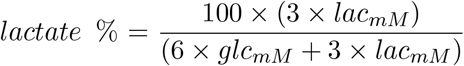. These results suggested non-trivial dynamics in the production of butyrate which could arise from competition for glucose and cross-feeding of L-lactate.

**Figure 5:**
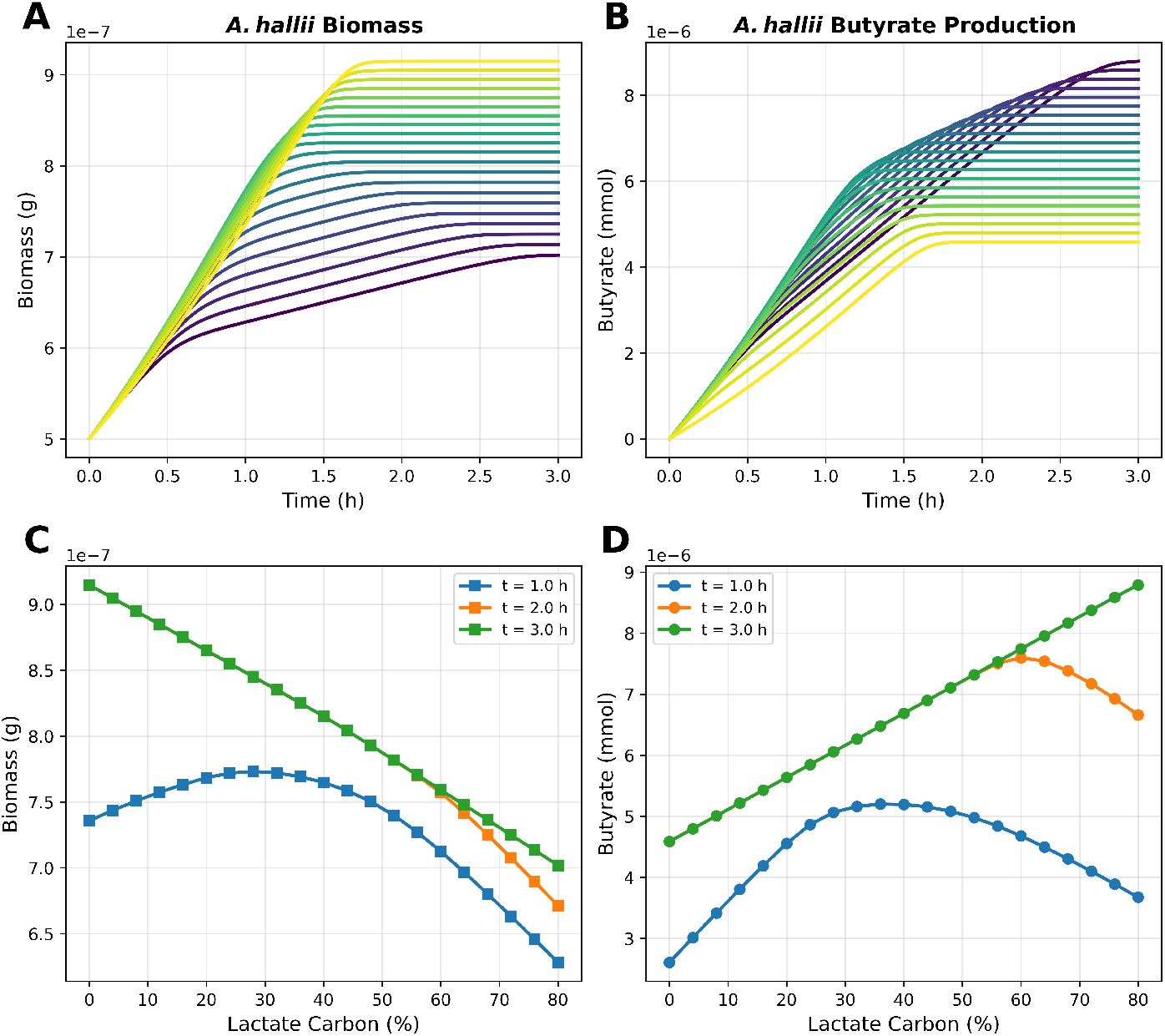
L-lactate carbon percentage sweep. (A) *A*.*hallii* biomass dynamics over time for various carbon compositions (increasing glucose: yellow, increasing L-lactate: blue). Increasing glucose leads to higher *A. hallii* final biomass. (B) Butyrate dynamics over time. Increasing L-lactate leads to higher final butyrate production. (C) *A. hallii* biomass vs. L-lactate carbon percentage at select time points. For finite time periods, an intermediate L-lactate percentage maximizes biomass, as shown at 1 h (blue line). (D) Butyrate vs. L-lactate carbon percentage at select time points. For finite time periods, an intermediate L-lactate percentage maximizes butyrate production, as shown at 2 h (orange line) and 1 h (blue line).

### 3.3 Spatio-Temporal Simulation in 2 Dimensions

This section provides the steps to run spatio-temporal simulations of *B. infantis* and *A. hallii* microbial community metabolism. A 2-dimensional spatial grid is defined to simulate the metabolism of two interacting colonies in a representative cross-section of the human colon outer mucus layer (3.3.1). The distance between these two interacting colonies is then varied to identify an optimal interaction distance for the production of butyrate (3.3.2).

#### 3.3.1 Mucus Layer

The goal of this section is to study how spatial structure influences competition and cross-feeding between *B*.*infantis* and *A*.*hallii*. Growth of both organisms is simulated on a 30 × 15 grid, which represent a 300 × 150 µm section of the human colon outer mucus layer. Glucose is supplied from the top row (lumen) at a static concentration of 10 mM. Both species are placed at row 2 (20 µm from the luminal surface) with initial biomass of 5 × 10^−10^ g. Bacterial colonies consume nutrients and produce metabolic by-products as they grow and spread horizontally across the mucus layer and vertically down towards the epithelial surface.

1. Define the Spatial grid representing the mucus layer cross-section. Set the total mucus depth to 150 µm. The space width and box volume are then defined based on this parameter. 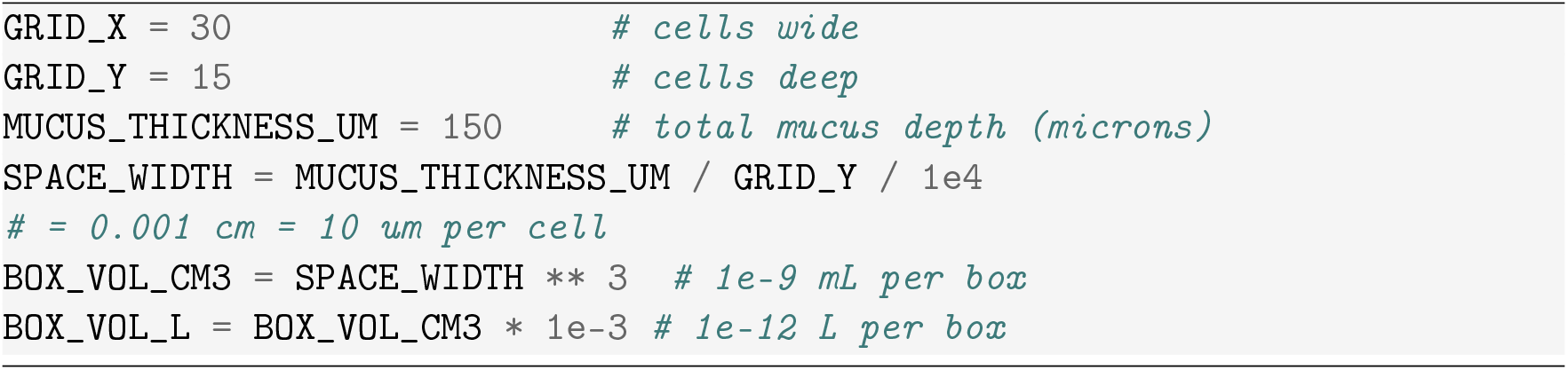
2. Define the diffusion coefficients for each metabolite. In our simulation, we used metabolite specific isotropic aqueous diffusion coefficients (Stewart, 2003). To account for the dense nature of the mucus, use a constant mucus reduction factor of 0.3 %. This parameter can be varied to change the density of the mucus. 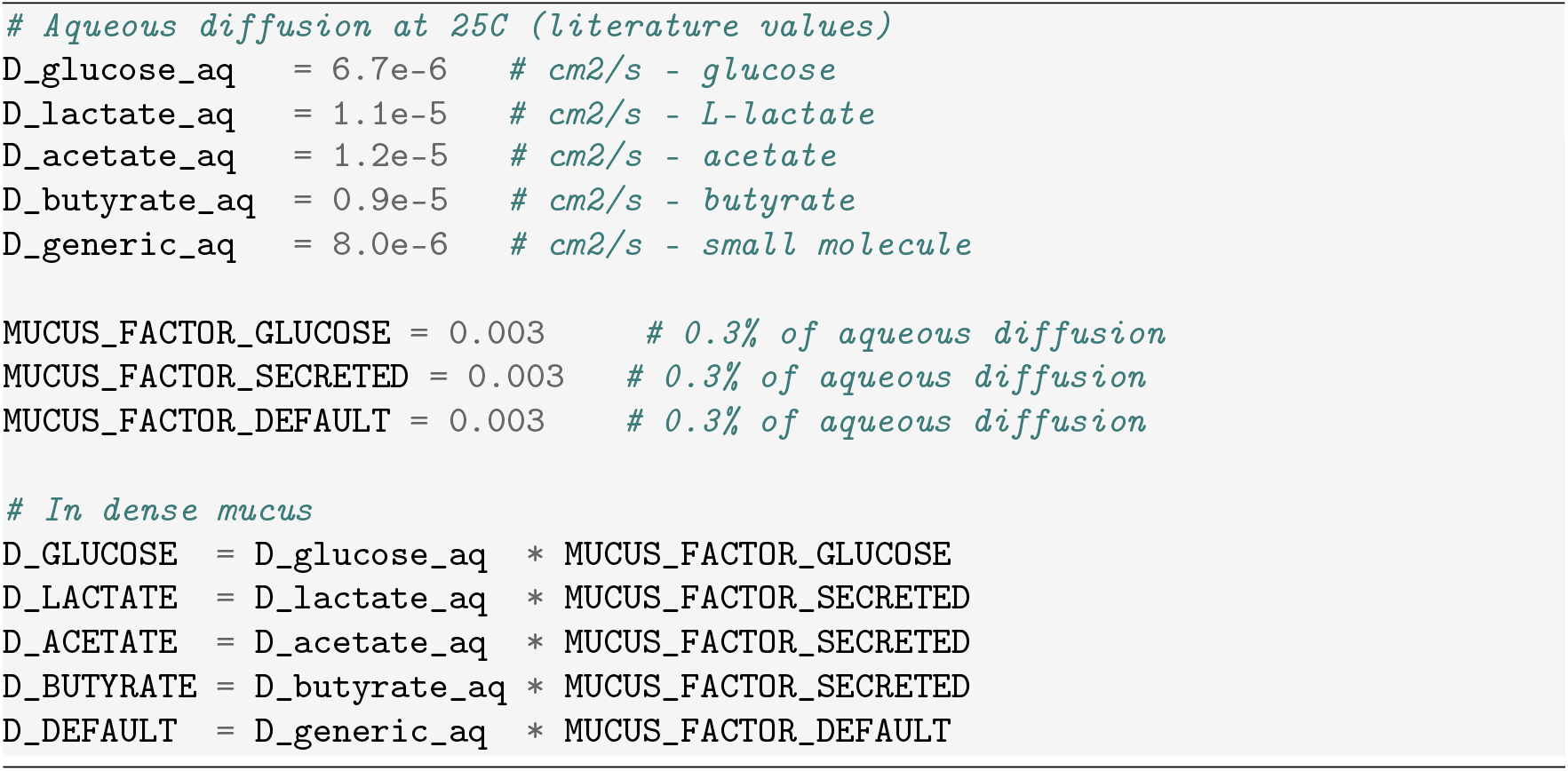
3. Set metabolite specific diffusion coefficients in COMETS layout. 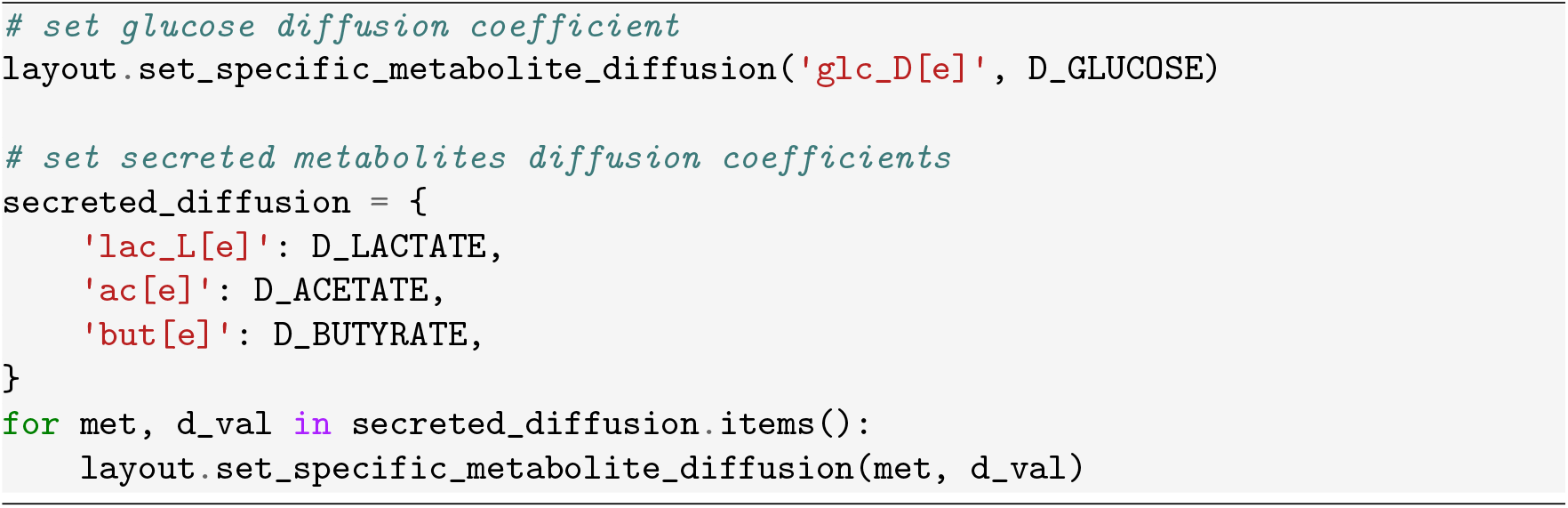
4. Set up the boundary conditions. Set a constant glucose concentration (10 mM) at the top boundary (*y* = 0, intestinal lumen) and zero glucose at the bottom boundary (*y* = 14, epithelial surface). 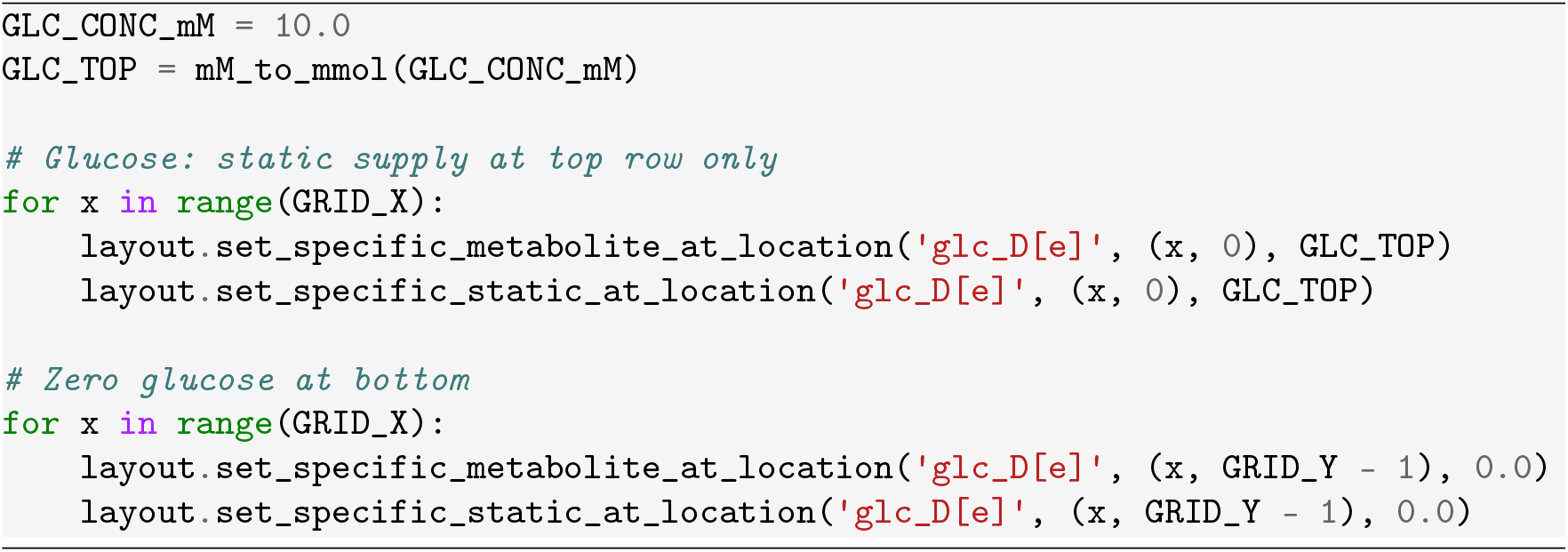
5. Place bacterial inocula at specific positions within the mucus layer grid. Place *B*.*infantis* at position (*x* = 7, *y* = 2) and *A*.*hallii* at position *x* = 22, *y* = 2 each with initial biomass of 5 × 10^−10^ g. 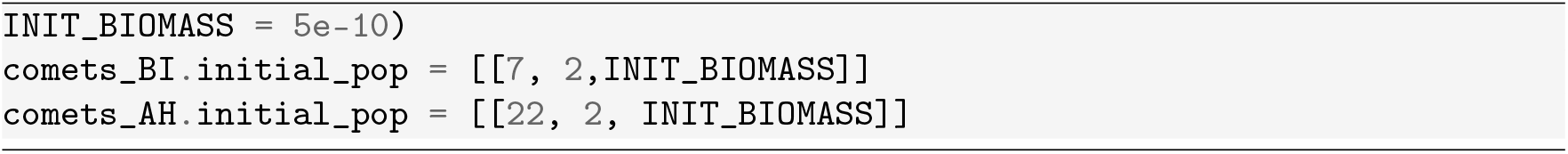
6. Configure nonlinear biomass diffusion. Set bacterial spreading to use density dependent nonlinear diffusion (*see* **Note 8**). 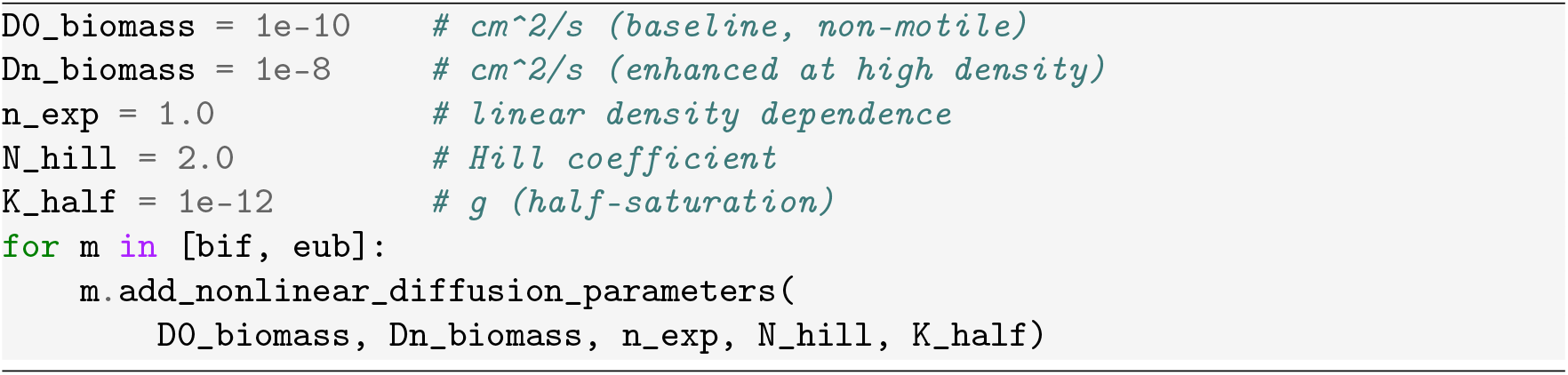
7. Set the COMETS parameters and run the simulation. This simulation takes around 2 minutes and 30 seconds to run. 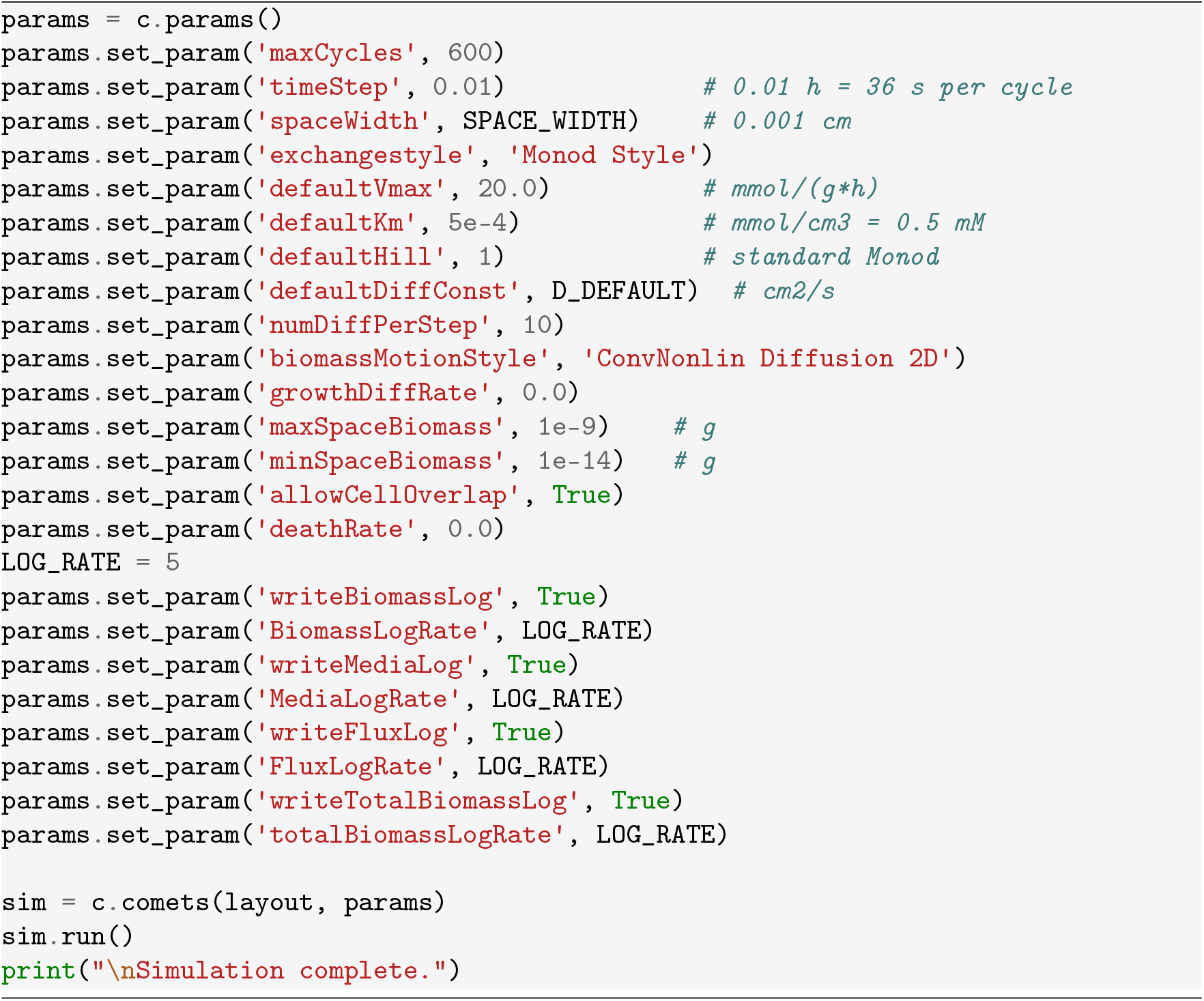
8. After the simulation completes, extract spatial data using the COMETS imaging functions. Apply “np.roll” to correct COMETS biomass indexing. (*See* **Note 9**). 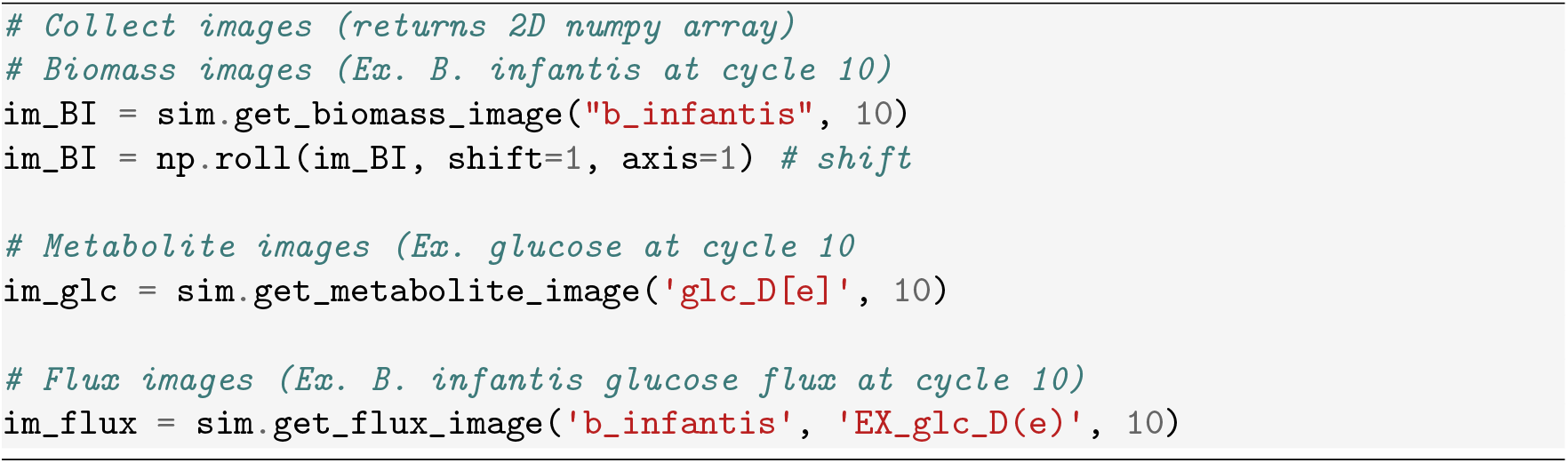
9. Plot biomass distributions. Examine the growth and spread of organisms by plotting their biomass at several time points (t = 0, 3, and 6 hours) as well as the total biomass over time (Figure 6).
10. Plot metabolite distributions to examine spatial and temporal dynamics. Plot the distributions of glucose, acetate, L-lactate, and butyrate at select time points (t = 0, 3, and 6 hours) as well as the total amounts of these metabolites over time (Figure 7).
11. Examine flux distributions. Plot biomass flux, indicating growth, for each organism along with key exchange fluxes for each organism (*B. infantis*: Glucose, L-lactate, and acetate; *A. hallii* : Glucose, L-lactate, and butyrate) (Figure 8).

**Figure 6:**
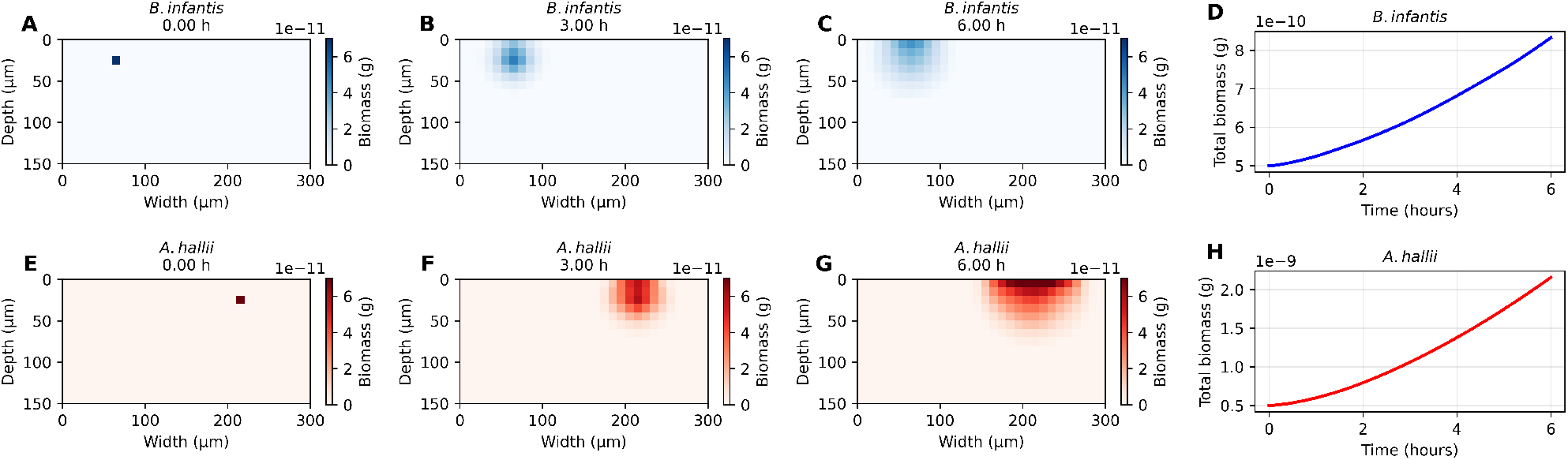
Spatial distribution of *B. infantis* and *A. halli* biomass at selected time points. Each organism spreads horizontally and vertically, establishing dense biomass colonies at the luminal (high-glucose surface) and growing down into the mucus (*B. infantis*: A-C and *A. halli* : E-G). The total biomass plots (*B. infantis*: D and *A. halli* : H) show that both organisms are growing, and *A. hallii* grows faster than *B. infantis*, consistent with previous results.

**Figure 7:**
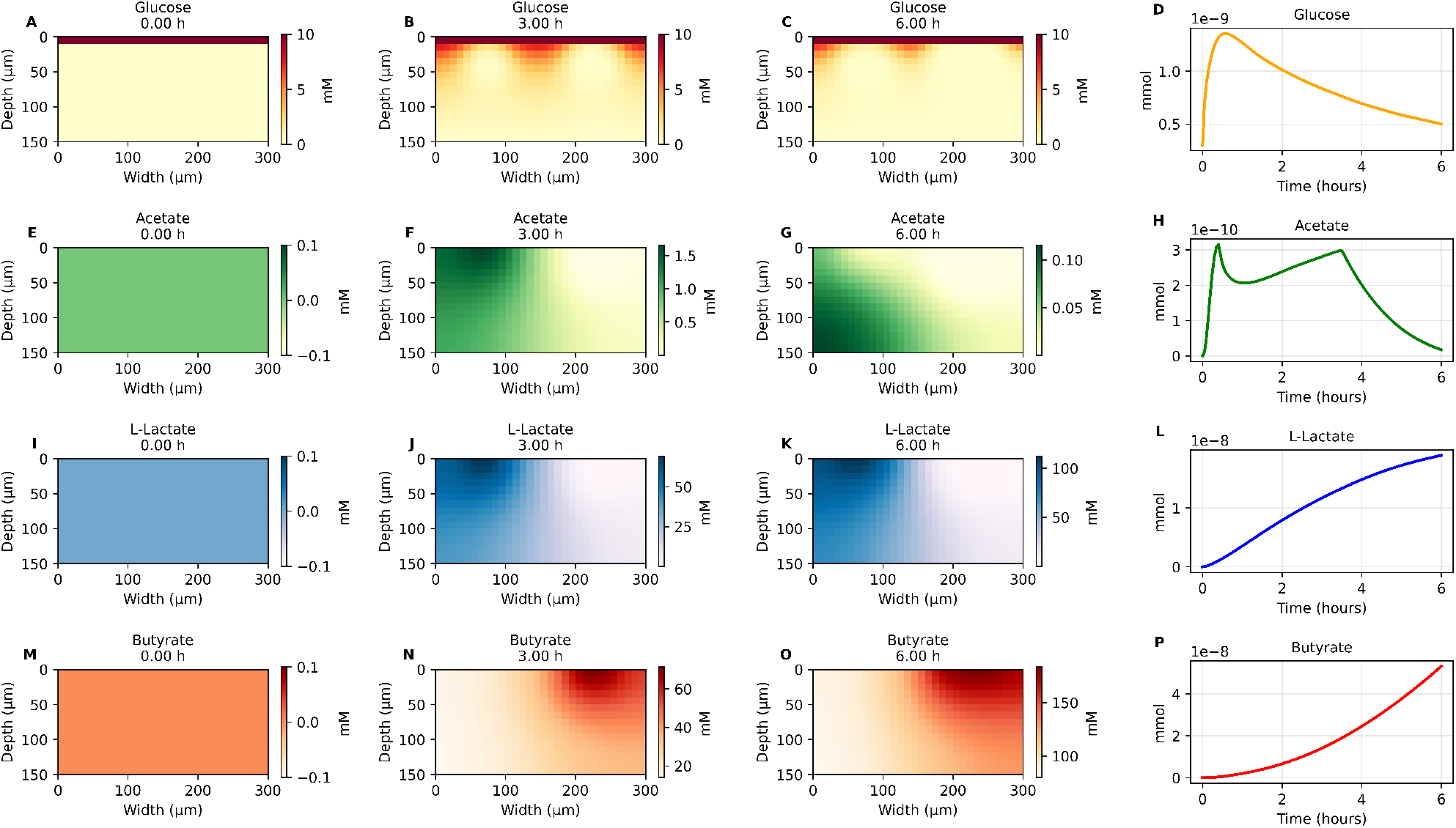
Spatial concentration distribution and total amounts of key metabolites: glucose, acetate, L-lactate, and butyrate. (A-D) Glucose initially increases as it diffuses into the system from the luminal source and is subsequently taken up by each organism and decreases over time as the organisms grow to higher biomass levels. (E-H) Acetate is sequentially produced and then consumed by each organism, and accumulates around *B. infantis* at 3 h before being consumed from the environment. (I-L) L-lactate increases around *B. infantis* and decreases around *A. hallii*. (M-P) Butyrate steadily increases around *A. hallii*. Note: metabolite colorbars do not have consistent limits across plots to enable visualization of spatial distributions.

**Figure 8:**
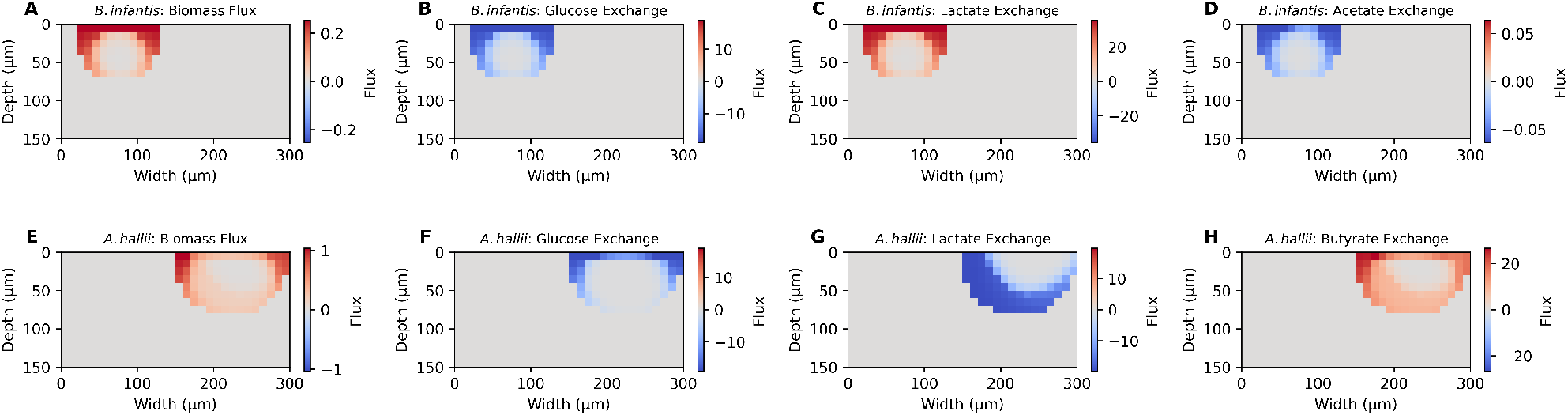
Spatial distribution of biomass and key exchange fluxes at final time point (6 hours). Both organisms grow at the periphery of the colony (A and E), where glucose is being actively taken up (B and F). *B. infantis* produces L-lactate and takes up acetate in symmetric patterns (C and D). *A. hallii* takes up L-lactate and produces butyrate with respective uptake and production asymmetrically greater on the side of the colony nearer to *B. infantis* (G and H). Blue color indicates uptake (negative exchange) while red indicates secretion (positive exchange). Biomass flux units are in h^−1^. Exchange flux units are in mmol/(g∗h).

#### 3.3.2 Effect of Species Distance on Butyrate Production

This section extends the spatial simulation to test how the distance between *B. infantis* and *A. halli* affects butyrate production. Butyrate production by *A. hallii* is fueled by glucose and L-lactate consumption. Close proximity to *B. infantis* decreases access to glucose (through competition) and increases access to L-lactate (through cross-feeding). Therefore, it is not clear how the distance between interacting colonies of *A. hallii* and *B. infantis* will affect butyrate production. We hypothesized that an intermediate distance would lead to an optimal production of butyrate and simulated different scenarios, initializing the colonies at various distances, to test this hypothesis.

**Table 2:** Spatial scenarios for investigating the effect of distance on butyrate production.

| Scenario | *B. infantis* Position ($x$) | *A. hallii* Position ($x$) | Separation ($\mu\text{m}$) |
| --- | --- | --- | --- |
| 1 | 14 | 16 | 20 |
| 2 | 12 | 18 | 60 |
| 3 | 10 | 20 | 100 |
| 4 | 8 | 22 | 140 |
| 5 | 6 | 24 | 180 |
| 6 | 4 | 26 | 220 |
| 7 | 2 | 28 | 260 |

1. Set up the spatial configurations varying only the horizontal positions of the two species. 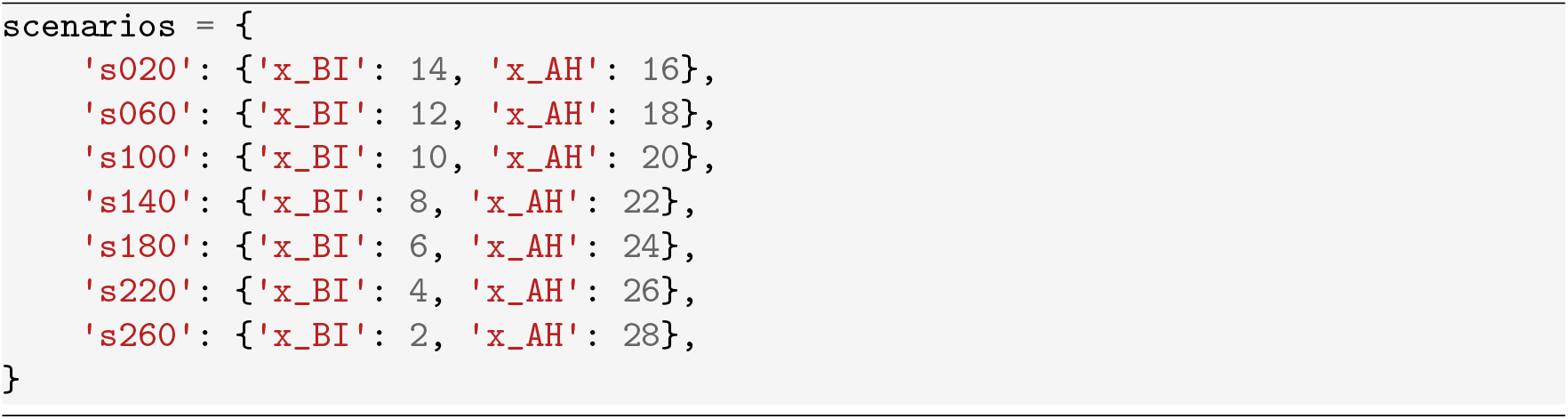
2. Create a run_scenario() function that includes simulation setup and accepts bacteria position as argument. These simulations follow the same procedure in section 3.3.1 with the only difference being the position of the two species. 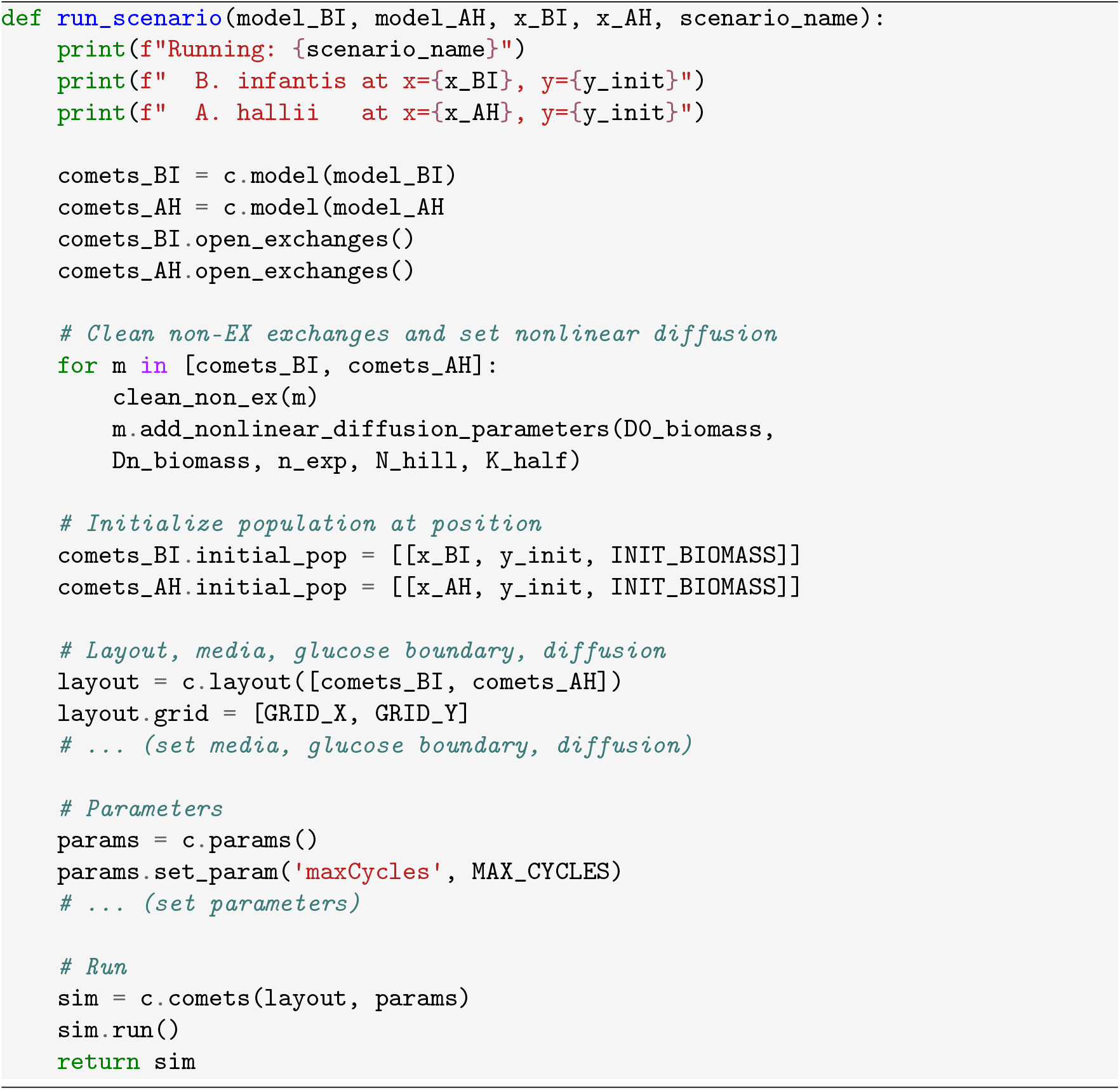
3. Run all scenarios. These simulations take around 17 minutes to run. 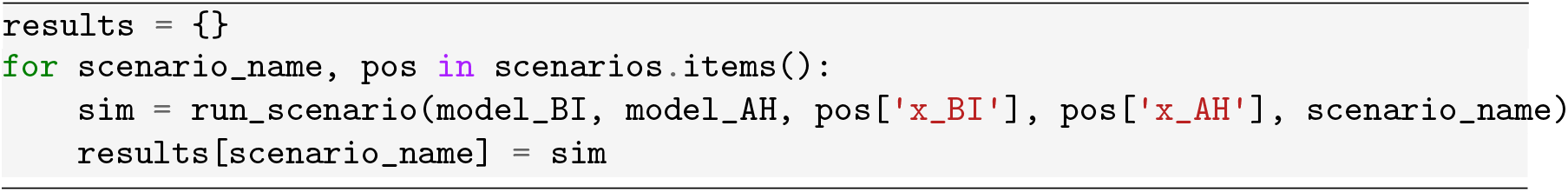
4. Compare total butyrate production across scenarios by plotting the resulting butyrate dynamics for each scenario (Figure 9).

**Figure 9:**
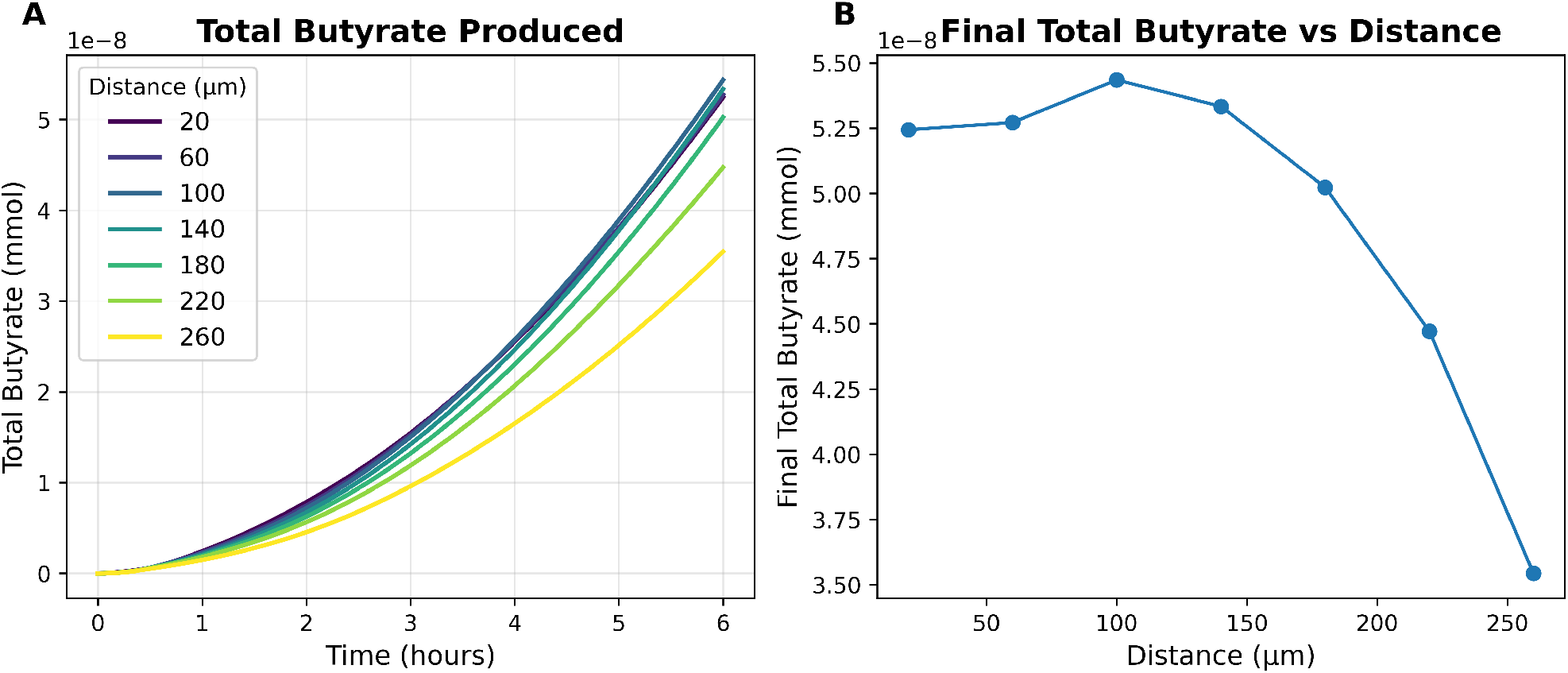
Variable distance simulation scenarios. (A) Butyrate dynamics for scenarios with initial distance between *B. infantis* and *A. halli* colonies ranging from 20 µm (blue) to 260 µm (yellow). (B) Final butyrate at 6 hours plotted against distance, demonstrating a maximum butyrate production at an intermediate distance of 100 µm.

## 4 Notes

1. AGORA2 models can be downloaded in either MATLAB (.mat) or SBML format. MATLAB models are loaded into COBRApy using the *cobra*.*io*.*load*_*matlab*_*model*() function. If you choose to use models in SBML format, then use the corresponding COBRApy I/O function *cobra*.*io*.*read*_*sbml*_*model*(). In our experience, the MATLAB models from the AGORA2 database are more reliable than the SBML models. So we choose to use these files when working with AGORA2.
2. AGORA2 models use database specific naming conventions. Examples are *EX*_*glc*_*D*(*e*) for the glucose exchange reaction and *glc*_*D*[*e*] for glucose. Additional information on these reactions and metabolites can be found on the Virtual Metabolic Human website (vmh.life). Different databases use different conventions, and it is important to match these identifiers exactly when setting the medium, boundary conditions, and diffusion coefficients.
3. Setting the COMETS path in the code is an optional step if you already have these paths configured in your system environment variables. If not, these variables need to be adjusted to your specific paths.
4. COMETS works internally with amount (mmol) per grid cell and not concentration. Hence, the need to define functions to convert to concentration for visualization and interpretation of results.
5. Additional information on COMETS parameters can be found in a table in the supplement of the COMETS Nature Protocols paper (Dukovski et al., 2021), starting on page 26.
6. COMETS sometimes flags internal reactions as exchange reactions. This can occur for sources and sinks, which are unbalanced internal reactions that are often included in COBRA models. To avoid this, we define a “clean_non_ex()” function that checks the model reactions and set EXCH = false for reactions that do not follow the EX_ naming convention. 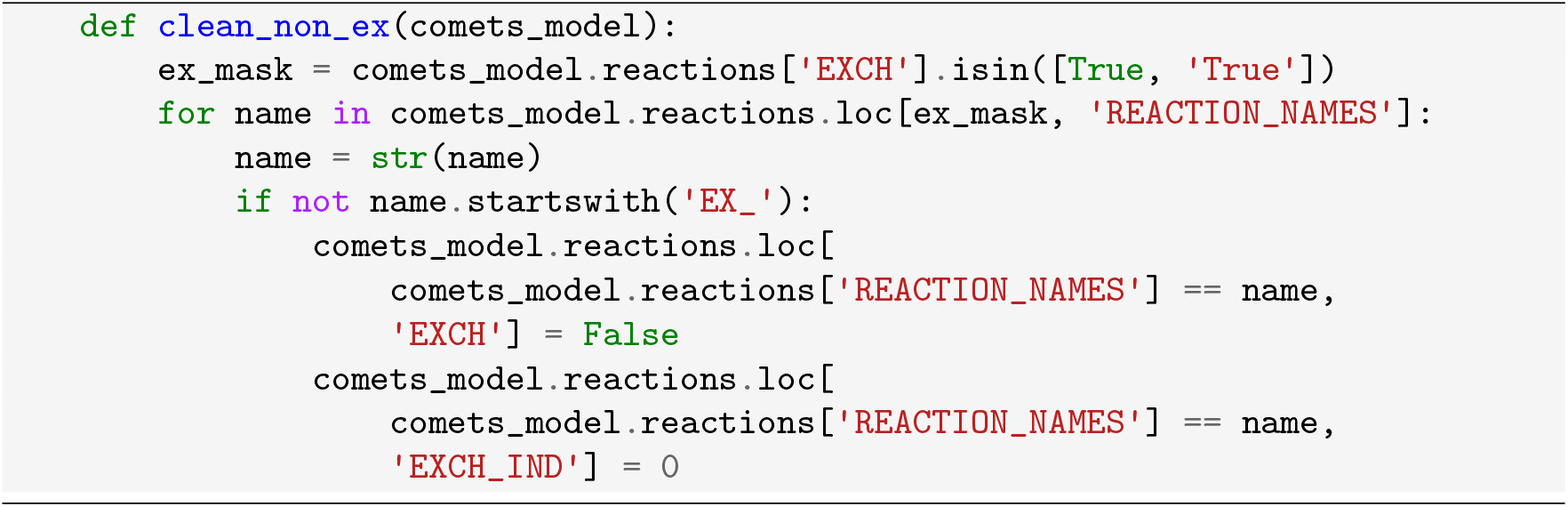
7. If model “A” has an exchange that model “B” lacks, COMETS may crash with an indexout-of-range error. Add closed dummy exchanges to resolve this. Build the union of all external metabolites from both models and add missing exchanges to each. 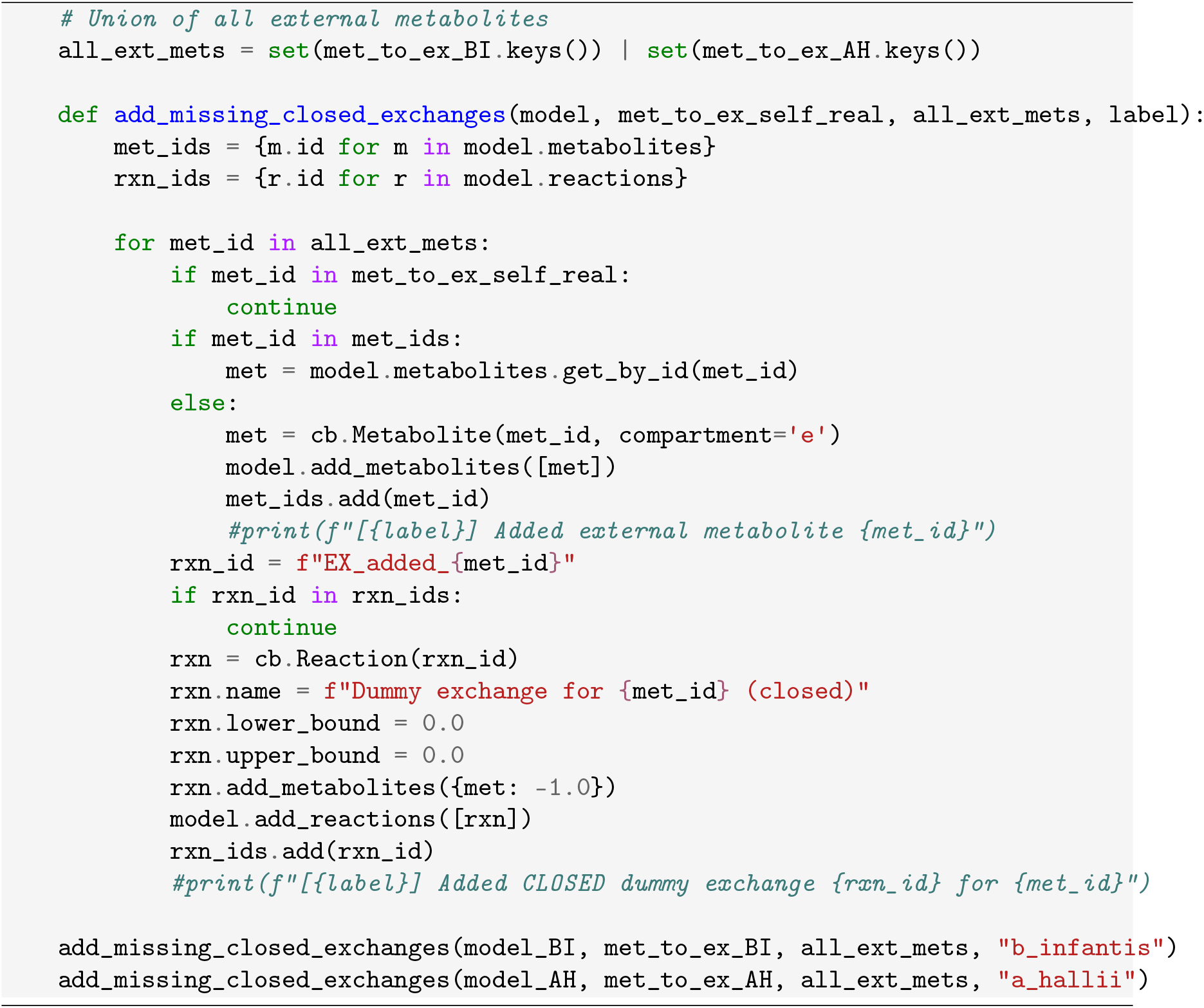
8. The biomass spread in the mucus layer is modeled using non-linear convective duffusion model (biomassMotionStyle = ‘ConvNonlin Diffusion 2D’). This model uses two parameters: a baseline diffusivity *D*_*o*_ = 10^−10^ cm^2^/s which represent non-motile bacteria in mucus and an enhanced diffusivity *D*_*k*_ = 10^−8^ cm^2^/s which represent mechanistic pushing at high local biomass density. A hill function with coefficient *k* = 2 and half saturation *K* = 10^−12^ g were used to modulate growth dependent diffusion. This approach limits the diffusion of non-growing biomass and is described further in the COMETS Nature Protocols paper supplement (Dukovski et al., 2021) and subsequent work (Dukovski et al., 2025).
9. The COMETSpy method “sim.get_biomass_image(species, cycle)” returns 2D array of biomass values indexed as (*x, y*). However, the returned array is shifted by one cell along the y-axis. Hence biomass that should appear at grid position *y* = *k* appear at *y* = *k* − 1. To fix this, we used the Numpy function “roll” to apply a shift of +1 along the y-axis. 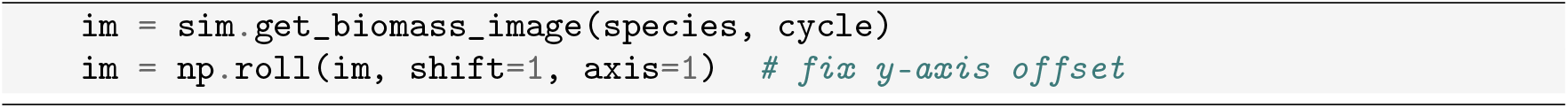

